# Spatial organization of transcribed eukaryotic genes

**DOI:** 10.1101/2020.05.20.106591

**Authors:** Susanne Leidescher, Johannes Ribisel, Simon Ullrich, Yana Feodorova, Erica Hildebrand, Sebastian Bultmann, Stephanie Link, Katharina Thanisch, Christopher Mulholland, Job Dekker, Heinrich Leonhardt, Leonid Mirny, Irina Solovei

**Author notes:** **MATERIALS & CORRESPONDENCE**: Irina Solovei.

## Abstract

Despite the well-established role of nuclear organization in gene expression regulation, little is known about the reverse: how transcription shapes the spatial organization of the genome. Owing to small sizes of most previously studied genes and the limited resolution of microscopy, the structure and spatial arrangement of a single transcribed gene are still poorly understood. Here, we make use of several long highly expressed genes and demonstrate that they form transcription loops with polymerases moving along the loops and carrying nascent RNAs. Transcription loops can span across microns resembling lampbrush loops and polytene puffs. Extension and shape of transcription loops suggest their intrinsic stiffness, which we attribute to decoration with multiple voluminous nascent RNPs. Our data contradict the model of transcription factories and indicate that although microscopically resolvable transcription loops are specific for long highly expressed genes, the mechanisms underlying their formation can represent a general aspect of eukaryotic transcription.

---

The understanding of eukaryotic gene transcription and the mechanisms of its regulation is progressively increasing at both the molecular^1-3^ and nuclear^4-6^ levels. Knowledge pertaining to an intermediate level of transcription organization, i.e. the spatial arrangement of a single expressed gene, is however surprisingly limited. Early works on gigantic chromosomes revealed that transcription units form loops emanating from a chromosome axis – so called lateral loops of lampbrush chromosomes^7^ and puffs of polytene chromosomes^8^. It was demonstrated that the 5’ and 3’ ends of loops are fixed in space and RNA polymerases II (RNAPIIs) move along a DNA template carrying a cargo of progressively growing nascent RNA transcripts (nRNAs)^7, 8^.

More recent studies of interphase nuclei have not revealed loops but identified transcription hubs consisting of clusters of expressed genes, aggregations of elongating RNAPIIs and nRNA accumulations^9-11^. These observations led to the popular hypothesis of *transcription factories*, which postulates that RNAPIIs are immobilized in groups, whereas activated genes of the same or different chromosomes are approaching a transcription factory and then reeled through immobilized RNAPIIs extruding nRNAs in a single spot^12, 13^.

The discrepancy between these two views on the mechanisms of transcription, can be explained to a great extent by the limitations of light microscopy resolution. Lateral lampbrush loops and polytene puffs are visible even under a phase contrast microscope^7, 8, 14^, owing to the significant length of transcription units (up to hundreds of kilobases) combined with their high transcriptional level. Therefore, for the visualization of expressed genes within interphase nuclei both a sufficient length and a sufficiently high expression are essential. The combination of these two traits, however, is rarely met in cultured mammalian cells, which are the major source of knowledge about transcription. Indeed, the majority of studied highly expressed genes are short^11^ and given that a 10 kb-long gene, when fully stretched, measures only 0.5 µm, it is comprehensible why their structure cannot be resolved by conventional microscopy with best possible resolution of 0.2-0.3 µm^15^. At the same time, long genes are generally not highly expressed, especially not in cultured cells^16, 17^.

To fill the gap in the knowledge about the spatial organization of transcription, we selected several genes that are both relatively long and highly expressed and studied their spatial arrangement in differentiated mouse cells. We demonstrated that these genes form microscopically resolvable transcription loops similar to lampbrush loops and polytene puffs. We provided evidence that transcription loops are decorated by elongating RNAPIIs moving along the gene axis and carrying nRNAs undergoing co-transcriptional splicing. Furthermore, we show that long highly expressed genes dynamically modify their harboring loci and extend into the nuclear interior presumably owing to their increased stiffness resulting from decoration with bulky nascent ribonucleoproteins (nRNPs). Collectively, our data indicate that although microscopically resolvable transcription loops are arguably specific for long highly expressed genes, the mechanisms underlying their formation can be general for eukaryotic transcription.

## RESULTS

### Selection of highly expressed long genes

For microscopic visualization, we searched for genes that are both relatively long and highly expressed, using thresholds for length of ≥100 kb, corresponding to the size of the smallest discernible lampbrush loops^18^, and for expression level of ≥ 1,000 TPM (transcripts per million), corresponding to the average expression level of the human GAPDH gene^19^.

Analysis of gene lengths and expression levels within the human and mouse genomes showed that less than 20% of genes are ≥100 kb (Extended Data Fig.1a) and that such genes, as a rule, are lowly expressed (Extended Data Fig.1b-d). Based on GTEx RNA-seq data, we selected 10 human genes that were above our thresholds and out of these further selected those that are expressed in cell types unambiguously identifiable in tissue sections (Supplementary Table1). The most highly expressed gene among them was the thyroglobulin gene (*TG; 268 kb; 7,510 TPM*) coding for the extracellular protein thyroglobulin secreted by thyrocytes; the other four selected genes encoded structural proteins of the contractile machineries of skeletal muscle (*TTG, NEB*) and smooth muscle (*CALD1, MYH11*) (Fig.1a).

**Figure 1.**
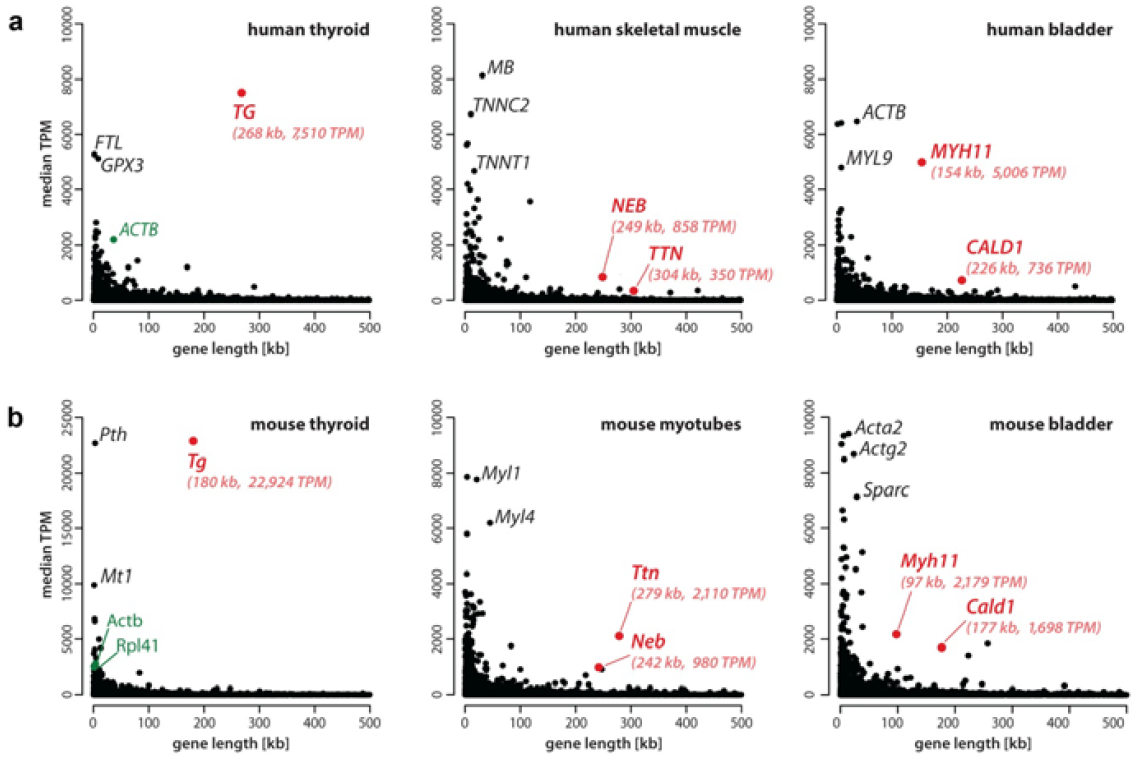
Selection of long highly expressed genes. **a**, Analysis of gene expression in selected human tissues (retrieved from the GTEx database. **b**, RNA-seq analysis of corresponding mouse tissues and cells from this study. Expression level (median TPM) is plotted against gene length according to GENCODE. Candidate genes with a length of ca. 100 kb or longer and an expression level of ca. 1,000 TPM or above are marked in *red*. Note the exceptionally high level of *Tg* expression, exceeding expression of housekeeping genes, such as *Actb* and *Rpl41* (marked in *green*). *Tg*, thyroglobulin; *Ttn*, titin; *Neb*, nebulin; *Myh11*, myosin heavy chain 11; *Cald1*, caldesmon 1; *Actb*, beta-actin; *Rpl41*, large ribosomal subunit protein EL41. For data on all protein coding genes see *Supplementary Tables 1* (human) and *2* (mouse).

Next, we performed RNA-seq analysis of mouse thyroid gland, myotubes, myoblasts and bladder tissue and confirmed the high expression of these five genes in the corresponding mouse cells (Supplementary Table2). In particular, the mouse thyroglobulin gene (*Tg*, 180 kb) is exceptionally upregulated (22,924 TPM) with an expression level 10-fold higher than that of some ubiquitously expressed housekeeping genes, such as non-muscle actin (*Actb*; 2,791 TPM) or ribosomal protein *Rpl41* (2,467 TPM) (Fig.1b). All five selected mouse genes – *Tg, Ttn, Neb, Cald1, Myh11* - met the conditions we have defined as prerequisites for our study (Fig.1b).

### Highly expressed long genes form microscopically resolvable Transcription Loops

Using genomic BAC probes encompassing the selected genes (Supplementary Table3), we carried out DNA-FISH on cryosections of the corresponding mouse tissues – thyroid gland (*Tg*), skeletal muscle (*Ttn* and *Neb*), heart muscle (*Ttn*), bladder and colon (*Myh11* and *Cald1*), as well as in cultured myoblasts (*Cald1*) and myotubes (*Ttn* and *Neb*). DNA-FISH, which includes RNase treatment and denaturation of cellular DNA, yielded two different signal patterns that were dependent on the expression status of genes. In the non-expressed state, genes were condensed into single compact foci sequestered to the nuclear periphery (Fig.2a). In expressed state, genes were strongly decondensed exhibiting several smaller foci in the nuclear interior (Fig.2b).

**Figure 2.**
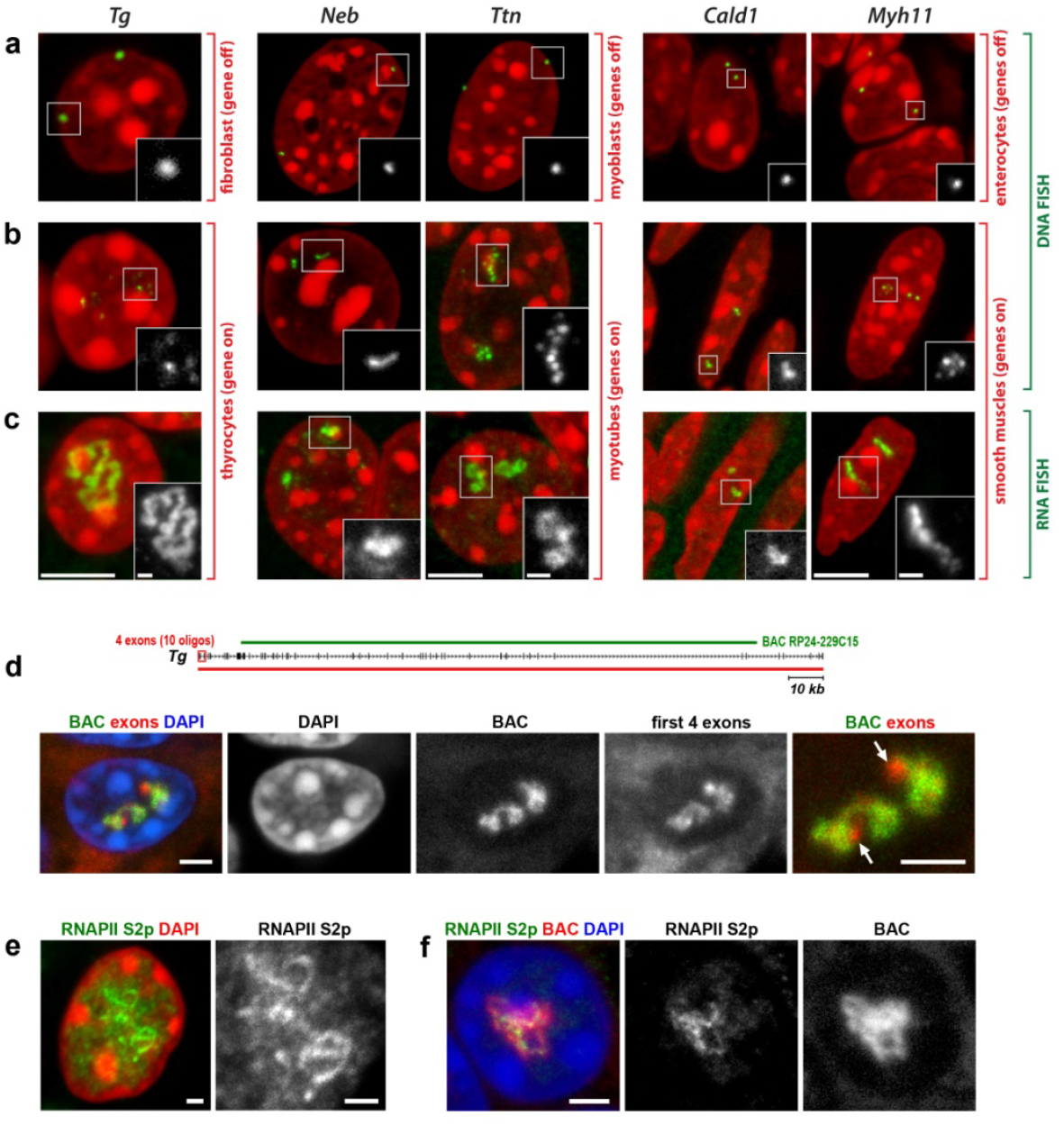
Highly expressed long genes form transcription loops (TLs). **a-c**, Five selected genes after either DNA-FISH (a,b) or RNA-FISH (c) with corresponding genomic probes. Control cells not expressing the respective genes exhibit focus-like condensed DNA signals at the nuclear periphery (a). In expressing cells, genes are strongly decondensed (b). RNA-FISH reveals TLs by hybridization to multiple nRNAs decorating the genes (c). **d**, An entire TL can be detected by oligoprobe hybridizing to 5’ exons, as exemplified for *Tg* TLs. The schematic shows distribution of the BAC covering mid-part of *Tg* (*green line*) and oligoprobe for the first 4 exons (*red rectangle*). Arrows point at *Tg* TL regions labeled by oligoprobe but not by BAC probe. **e**, *Tg* TLs revealed by immunostaining of elongating RNAPII (Ser2p). **f**, Immuno-FISH showing colocalization of structures marked with elongating RNAPII and *Tg* TLs. Images are *projections of 1-2*.*5 µm* confocal stacks. Scale bars: a-c, *5 µm*, in insertions, *1 µm*; d, f, *2 µm*; e, *1 µm*.

Next, we performed RNA-FISH on the same set of samples, omitting both RNase treatment and cell DNA denaturation. With this setup, the same genomic probes used for DNA-FISH hybridize to nRNAs. As expected, in cells not expressing the genes, FISH signals were absent, whereas expressing cells exhibited massive RNA signals (Fig.2c, Extended Data Fig.2). The RNA-FISH signals were extended and either had a shape of a coiled loop (e.g., *Tg, Ttn*), or formed less discernible elongated structures (e.g., *Neb, Cald1*). The loops were particularly prominent in case of the *Tg* gene - they spread throughout the nuclear interior and measured up to 10 *µm* (Extended Data Fig.2a; Supplementary video1).

Since introns, as a rule, are substantially longer than exons, we assumed that the used genomic probes mostly hybridized to unspliced introns of nRNAs, thereby outlining the contours of transcribed genes. The entire expressed gene, however, can also be visualized by using oligoprobes hybridizing to several 5’ exons (Fig.2d). In agreement with FISH data, immunostaining of elongating RNAPII in thyrocytes revealed strongly convoluted structures colocalizing with RNA-FISH signals and apparently corresponding to the *Tg* gene axis covered by polymerases (Fig.2e,f). Hereafter we refer to these structures as *Transcription Loops (TLs)*.

TL signals varied in shape and coiling degree and no consistent pattern of loop folding or location was observed with exception to their invariably interior positioning. Scoring of RNA-FISH signals revealed that all five selected genes are expressed mostly biallelically (Extended Data Fig.3), which is in line with the observations that monoallelic expression^20^ is more common for lowly expressed genes^21^.

At the light microscopy level one can trace only the general contour of the TLs, but within this contour there is apparently finer coiling of the gene axis not resolvable by deconvolution or high-resolution microscopy (Extended Data Fig.4a). To demonstrate the inner structure of TLs, we performed FISH with the *Tg* probe on thin thyroid sections (50-70 nm) and indeed observed such coiling (Extended Data Fig.4b). Therefore, the compaction level of *Tg* TLs compatible with nucleosomal chromatin (ca. 17 kb/µm), which was calculated based on contour length measurements (Extended Data Fig.4c), is rather an overestimation. Hence, we cannot exclude that during *Tg* transcription a significant proportion of nucleosomes is lost, similarly to ribosomal genes^22^ or lampbrush chromosome loops^23^.

### Transcription Loops manifest the progression of transcription and co-transcriptional splicing

To confirm that the observed RNA signals are not accumulations of messenger RNAs (mRNAs) but represent nRNAs, we performed two types of RNA-FISH experiments. Firstly, we used differentially labeled probes hybridizing to 5’ exons (2-12) and 3’ exons (33-47) of the *Tg* gene. We reasoned that if loops represent mere mRNA accumulations, both probes would label the entire structures. The probe for 3’ exons, however, labeled only the second half of TLs (Fig.3a), confirming that TLs are decorated with growing nRNAs. On a smaller scale, we demonstrated that oligoprobe for 5’ half of *Tg* intron 41 labels the entire intron, whereas oligoprobe for 3’ half of the intron labels only half of it (Fig.3b).

**Figure 3.**
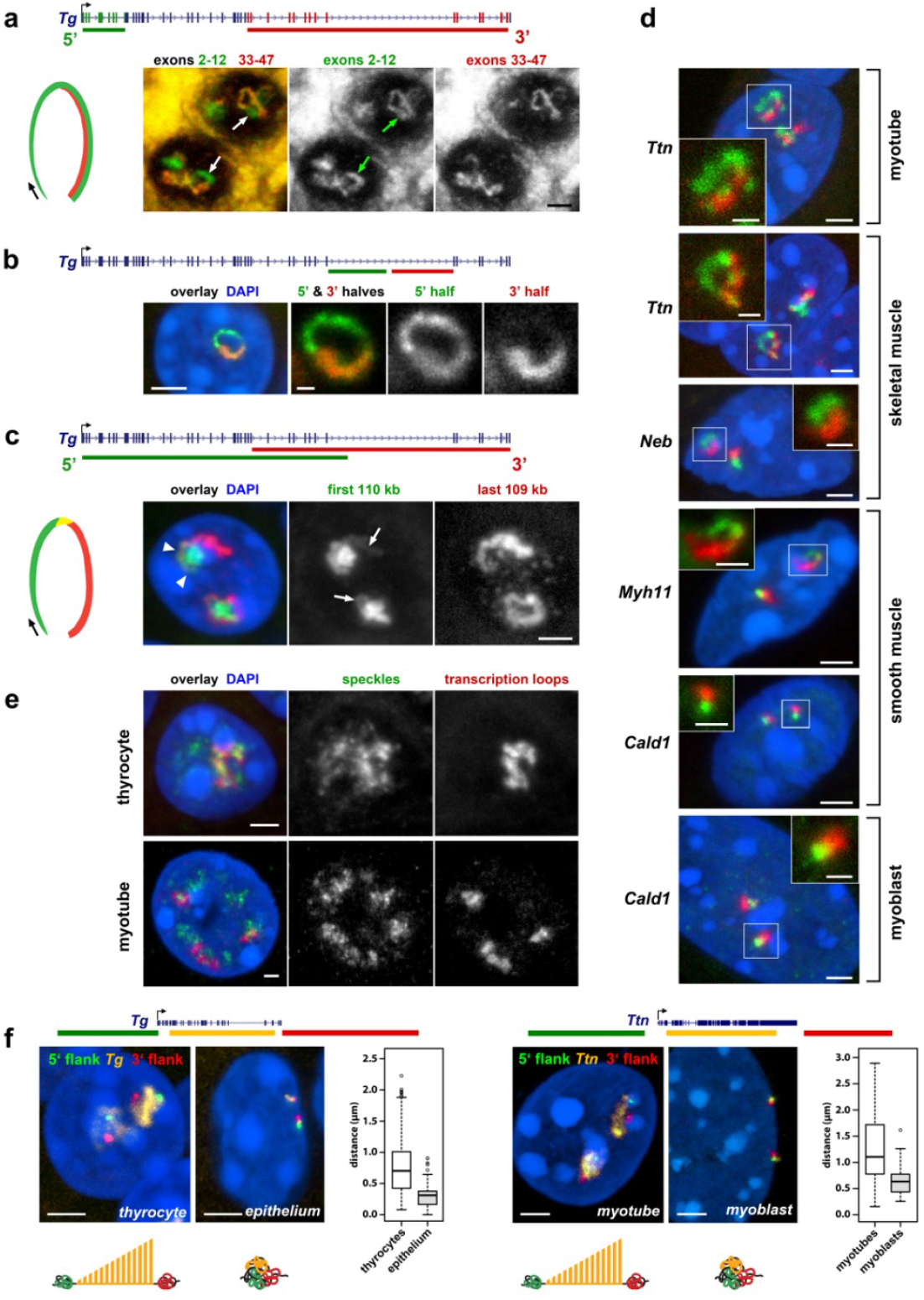
**TLs manifest transcription progression and dynamically modify harboring chromosomal loci.** **a**, Successive labeling of *Tg* TLs with probes for 5’ (*green*) and 3’ (*red*) exons. Cartoon on the *left* shows changes in the exon composition of nRNAs during transcription progression; examples of *Tg* TLs labeling after RNA-FISH are on the *right. Arrows* point at the TL regions labeled with only 5’ probe. **b**, Successive labeling of *Tg* long intron 41 with oligoprobes for the entire intron (*green*) and for its second 3’ half (*red*). **c**, Sequential labeling of *Tg* TLs with genomic probes highlighting 5’ (*green*) and 3’ (*red*) introns. Cartoon on the *left* shows changes in the intron composition of nRNAs during transcription progression; example of *Tg* TLs labeling after RNA-FISH is shown on the *right*. Genomic probes highlighting 5’ and 3’ introns sequentially label the TLs as a result of co-transcriptional splicing. Since the 5’ probe includes 5’ exons, it also faintly labels the second half of the loop (*arrows*). The region hybridized by both overlapping BACs is marked with *arrowheads*. **d**, TLs formed by other long highly expressed genes exhibit co-transcriptional splicing. RNA-FISH with probes encompassing 5’ (*green*) and 3’ (*red*) regions of the genes. **e**, Immuno-FISH visualizing nuclear speckles (SON staining) and TLs of *Tg* (*left*) and *Ttn* (*right*). Note that the myotube nucleus is tetraploid (4 *Ttn* signals). **f**, Expressed genes expand from the harboring loci and separate their flanks. Distances between 5’ (*green*) and 3’ (*red*) flanking regions of the *Tg* and *Ttn* genes are larger in cells expressing the genes compared to control cells with silent genes (*left and right panels, respectively*), as reflected in corresponding cartoons at the bottom. The boxplots depict the 3D distance between the flanking regions in expressing and not-expressing cells. The median inter-flank distance for *Tg* in thyrocytes is 2.3-fold larger than in control epithelial cells (703 nm *versus* 311 nm). The median inter-flank distance for *Ttn* in myotubes is 1.7-fold larger than in control myoblasts (1,104 nm *versus* 634 nm). Projections of confocal sections through 1 - 3 *µm*. Scale bars, *2 µm*, in insertions on *d, 1 µm*. Distributions of used probes in respect to the studied genes are depicted above the image panels.

Secondly, we used genomic probes highlighting 5’ and 3’ introns. We expected that the two probes will hybridize to nRNAs on TLs in a consecutive manner as a result of co-transcriptional splicing and, indeed, observed such a pattern. The 5’ genomic probe strongly labeled the first half of the loop by hybridizing to nRNAs containing yet unspliced 5’ introns and the 3’ probe labeled the second half of the loop by hybridizing to yet unspliced 3’ introns (Fig.3c,d). The oligoprobes covering 1-5 kb of two sequentially positioned introns of *Tg, Ttn* and *Cald1* labeled TLs only partially, producing separate and non-overlapping signals (Expanded Data Fig.5), further confirming the ongoing splicing^24^. In agreement with multiple splicing events occurring along highly expressed genes, TLs are either closely adjacent to or even colocalize with nuclear speckles (Fig.3e).

Besides TLs, RNA-FISH with genomic probes revealed numerous granular signals scattered throughout the nucleoplasm, likely representing both mRNA and accumulations of spliced out but not yet degraded introns (Extended Data Fig.6a). In myotubes, the majority of granules (81%) are double-labeled with probes for 5’ and 3’ halves of *Ttn* and found in both the nucleoplasm and cytoplasm and thus apparently represent *Ttn* mRNAs. Remarkably, the two signals are spatially distinguished within the granules presumably due to the exceptionally long *Ttn* mRNA of ca 102 kb (Extended Data Fig.6b). mRNAs of *Tg, Neb, Cald1* and *Myh11* genes are short (4 - 20 kb) and can be only detected with, e.g., oligoprobes specifically hybridizing to all exons (Extended Data Fig.6c). Therefore, in these cases, genomic probes detect accumulations of excised introns (Extended Data Fig.6d).

### Transcription Loops are open loops with separated flanks

It is widely believed that physical interactions between the TSS and TTS are crucial for a high level of expression. Such interactions have been interpreted as a ‘bridge’ for RNAPIIs enabling them to immediately reinitiate transcription after termination, i.e. to transcribe “in circles”^25^. To assess whether the beginnings and the ends of the selected long highly expressed genes reside in close proximity to each other, we visualized TLs and their genomic flanks for *Tg, Ttn* and *Myh11* genes and found that the 5’ and 3’ gene flanks are visibly separated in 85%, 87% and 90% of the respective alleles. In particular, 3D distance measurements between flanks showed that they can be separated by up to 2.2 *µm* (*Tg*) and 2.9 *µm* (*Ttn*), and that the inter-flank median distances in expressing cells are ca. 2-fold larger compared to control cells not expressing these genes (Fig.3f). This finding demonstrates that, contrary to the “transcription cycle” hypothesis, TLs of long highly expressed genes are open loops with separated flanks.

### Transcription Loop size is not proportional to gene length but to gene expression

Gene length alone is not a good predictor of TL loop size but rather a high expression level is required: e.g., RNA-FISH signals of the longer *Ttn* gene (279 kb; 2,110 TPM) are less expanded compared to those of the shorter but 10-fold more highly expressed *Tg* gene (180 kb; 22,924 TPM). Notably, the size of RNA-FISH signals inversely correlates with condensation of genes detected by DNA-FISH: e.g., the gene body of *Tg* is strongly decondensed and marked with only few small faintly labeled foci, whereas the *Ttn* gene is more condensed and displays a chain of several larger foci with small gaps in between (Fig. 4a,b). The *Neb* gene, whose expression is more than 2-fold lower than that of *Ttn*, typically exhibits even more prominent gene body condensation. The body of *Dmd*, the longest mammalian gene (ca. 2.3 Mb) with a very low expression (5 TPM), remains condensed (Fig. 4c,d).

**Figure 4.**
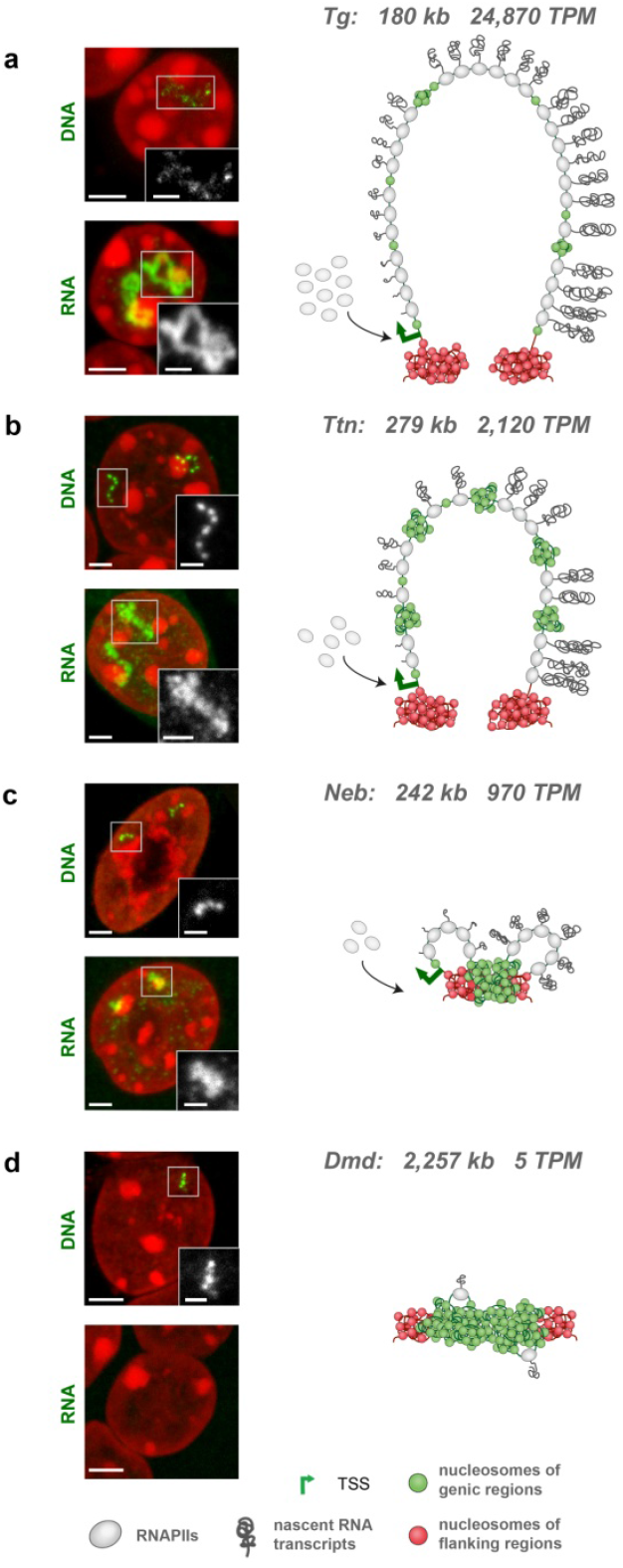
**TL size is disproportional to gene length but correlates with gene expression.** Examples of 4 long genes - *Tg* (**a**), *Ttn* (**b**), *Neb* (**c**) and *Dmd* (**d**) - arranged from top to bottom according to their expression level in the respective cell type. Gene length and expression level are indicated next to a gene symbol. For every gene two images are displayed *on the left*, showing DNA-FISH detecting the gene body and RNA-FISH detecting nRNAs. The more a gene is expressed, the less solid the DNA-signal is and the more expanded the RNA-signal is. *Vice versa*, the less a gene is expressed, the more condensed the gene body is and the less extended the RNA-signal is. The schematics on the right are speculative interpretations of the observed FISH signal patterns in terms of transcriptional bursts, depicted as RNAPII convoys with attached nRNAs, and transcriptional pauses, depicted as condensed chromatin (*green nucleosomes*). For simplicity, splicing events are not shown on the schemes. Microscopic images are projections of 1-1.5 µm confocal stacks. Scale bars, *2 µm*, in insertions, *1 µm*.

The coordinated pattern of gene body condensation and TL expansion presumably reflects the frequency and duration of transcriptional bursts that are directly related to the expression level of genes^26^. The possible scenario is that a highly expressed gene is transcribed in long and frequent bursts; an intermediately expressed gene – in short, less frequent bursts; the bursts of a lowly expressed gene are even shorter and less regular (Fig. 4). Consequently, in case of high expression, a gene is covered by long and frequent RNAPII convoys^27^ and the entire gene body forms a transcription loop (e.g. *Tg, Ttn*). In case of lower expression, short and infrequent bursts do not decondense the gene body entirely but rather form short and sparse RNPII convoys resulting in small “travelling” transcription loops (e.g., *Neb, Cald1*). In both cases, however, transcription occurs via the same mechanism with RNAPIIs moving along a gene, which is evident through sequential highlighting of introns (Fig. 3b,c).

### Transcription Loops are dynamic structures

To confirm that TLs are formed upon transcriptional activation, we performed several experiments to activate or inhibit *Ttn* transcription *in vitro*. First, we aimed to induce *Ttn* TLs in cells that do not express this gene and generated myoblasts stably expressing dCas9 conjugated with the tripartite transcription activator VP64-p65-Rta (VPR)^28^. Targeting the dCas9-coupled activator to the *Ttn* promoter region induced expression of *Ttn* in myoblasts to a level comparable to that in myotubes, which was accompanied by formation of *Ttn* TLs and by a 2-fold increase of *Ttn* median inter-flank distances (Fig.5a).

**Figure 5.**
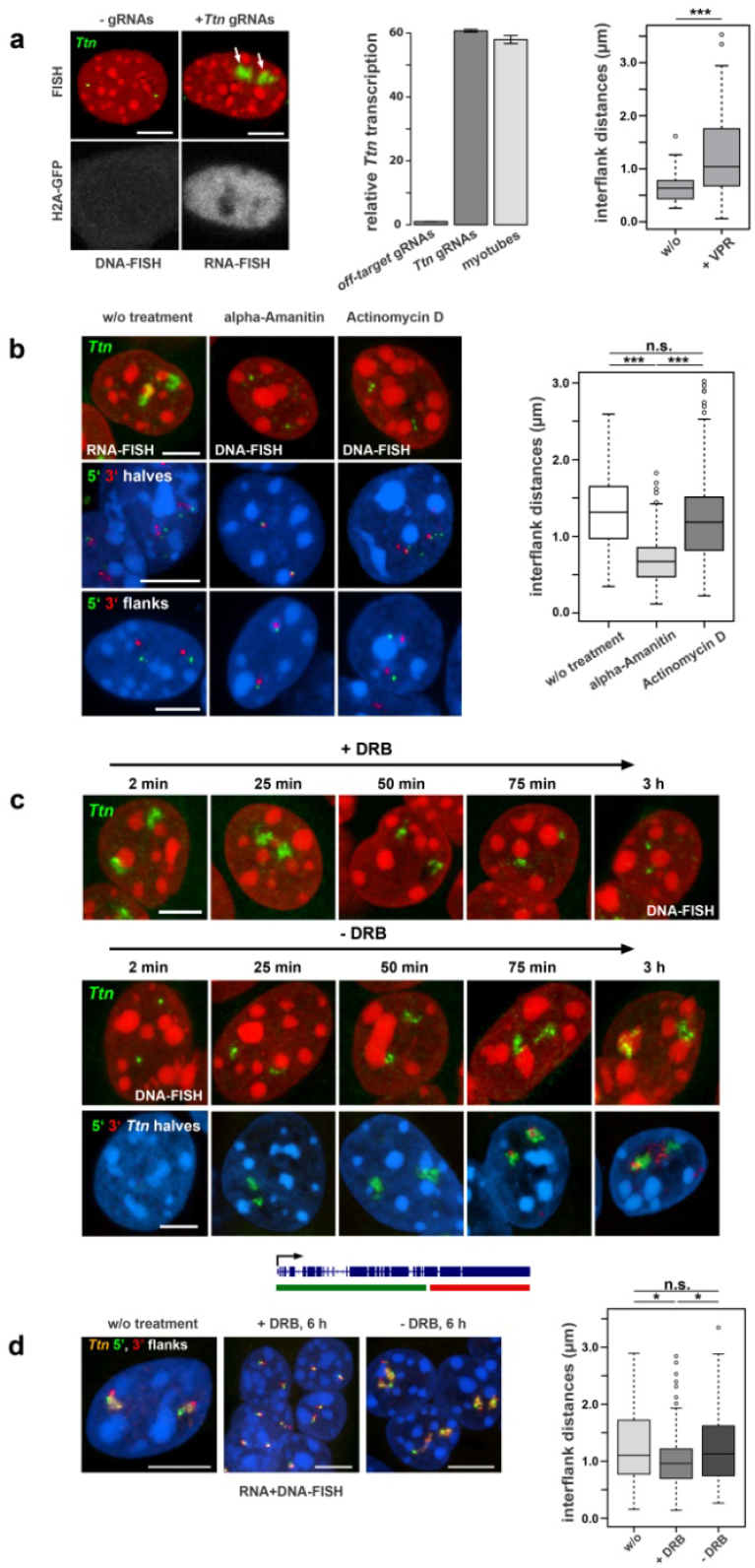
TLs are dynamic structures. **a**, Ectopic induction of *Ttn* TLs in myoblasts not expressing *Ttn*. Myoblasts, stably expressing dCas9-VPR, were co-transfected with plasmids for H2A-GFP and with either gRNAs targeting the *Ttn* promoter or with off-target gRNAs. As assessed by qPCR, *Ttn* expression was induced in cells transfected with *Ttn* gRNAs to a level comparable to that of untreated myotubes (*error bars depict standard deviations)*. Accordingly, 90% of transfected myoblasts exhibited *Ttn* TLs and a 2-fold increase in median inter-flank *Ttn* distances (from 634 nm to 1,039 nm) *(***p-value < 0*.*001, Wilcoxon rank sum test)*. In untransfected cells, the *Ttn* gene remained condensed with converged flanks. **b**, Treatment with transcription inhibitors eliminates *Ttn* TLs in differentiated myotubes. *Ttn* TLs are not detectable after treating cells with α-amanitin (*mid column*) and actinomycin D (*right column*). According to their different mechanisms of transcription inhibition, the two drugs affect gene condensation in different ways. As DNA-FISH with probes for 5’ and 3’ halves (*mid row*) and 5’ and 3’ flanks (*bottom row*) shows, α-amanitin treatment caused strong condensation of the gene body and convergence of the flanks, whereas after actinomycin D treatment, the gene body remains decondensed and flanks remain diverged, similar to *Ttn* in untreated cells (*left column*). (*\*\*\*p-value < 0*.*001, n*.*s. = not significant, Wilcoxon rank sum test*). **c**, Application of DRB, a reversible transcription inhibitor, causes inhibition of transcription elongation and thus leads to gradual shrinkage of *Ttn* TLs (*upper panel*); removal of DRB revives elongation and leads to a gradual restoration of *Ttn* TLs (*mid panel*). Differential labeling of 5’ (*green*) and 3’ (*red*) halves of *Ttn* (*see gene labeling scheme at the bottom*) allows better monitoring the TLs dynamics: the signal for the 5’ end appears first, the signal for 3’ end appears after ca. 1 h (*bottom panel*). **d**, *Ttn* inter-flank distances are decreased after complete transcription inhibition with DRB from 1,104 nm to 963 nm, but remain larger than inter-flank distances in myoblasts (634 nm); upon transcription restoration the distances are restored up to 1130 nm (*\*p-value < 0*.*05, n*.*s. = not significant, Wilcoxon rank sum test)*. For used *pseudo-colors* and more inter-flank distances of *Ttn* see Figure 3f. Scale bars, *5 μm*.

Conversely, we aimed to eliminate *Ttn* TLs in differentiated myotubes by transcription inhibition. To this end, we incubated myotubes with α-amanitin or actinomycin D and demonstrated that both drugs cause abortion of RNAPIIs and disappearance of TLs (Fig.5b). In agreement with their different mechanisms of action^29^, the drugs differently affected gene body condensation. In case of α-amanitin, which prevents DNA and RNA translocation by binding near the catalytic site of RNAPII, the *Ttn* gene condensed and the gene flanks converged. In case of actinomycin D, which prevents RNAPII progression by intercalating into the DNA minor groove, *Ttn* remained decondensed with separated flanks (Fig.5b).

To monitor TL dynamics further, we treated differentiated myotubes with 5,6-Dichloro-1-β-D-ribofuranosylbenzimidazole (DRB), a drug that reversibly prevents transcription elongation^29^. As anticipated, during the first 2 hours of drug treatment, nRNA signals diminished and the gene body condensed. After drug removal, the TL signal was gradually restored with the 5’ signal emerging first and the 3’ signal emerging only after ca. 1 h (Fig.5c). In accordance with transcriptional changes, *Ttn* flanks converged upon drug application and diverged after drug removal (Fig.5d).

### Transcription Loops of long genes can expand from harboring chromosomes and cause local chromosomal reorganization

Chromosome territoriality is one of a central doctrine of nuclear organization^30^. To investigate the relationship between TLs and their harboring chromosomes, we visualized the *Tg* TL and chromosome 15 by FISH detecting both DNA and RNA. We found that the *Tg* TL is excluded from the chromosome territory and either expands into the nucleoplasm or coils next to the chromosome, forming its own “transcription territory” (Fig.6a). The areas of TLs are often depleted of chromatin, indicating an extremely high concentration of proteins involved in the massive *Tg* transcription (Fig.6b). Remarkably, in 2% of *Tg* alleles, TLs split chromosome 15 territories into two halves with a gap marked with the 5’ and 3’ *Tg* flanking sequences and filled with *Tg* TLs (Fig.6c,d).

**Figure 6.**
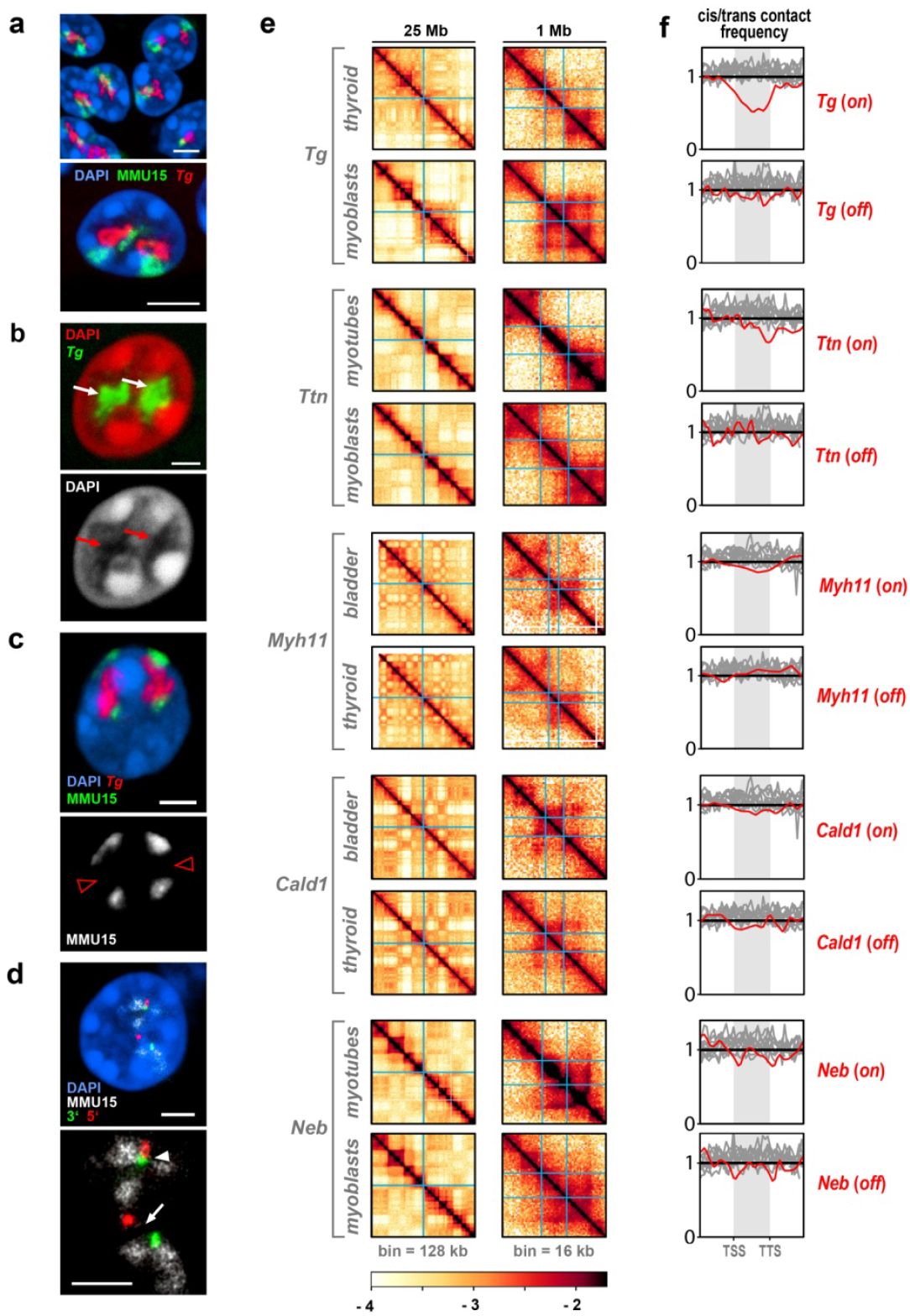
TLs are excluded from harboring chromosomes. **a**, *Tg* TLs (*red*) emanate from harboring chromosome 15 (*green*) and protrude into the nuclear interior. **b**, Nucleoplasmic regions occupied by *Tg* TLs (*white arrows*) are depleted of DAPI-stained chromatin (*red arrows*). **c**, Ca. 2% of *Tg* alleles (*red*) split chromosome territories (*green*) into two halves with an unpainted gap between them (*red empty arrowheads in the lower panel*). **d**, The gap in the chromosome territory (*arrow*) is marked with the 5’ (*red*) and 3’ (*green*) *Tg* flanking sequences. In the second homologue chromosome, the chromosome is not split and the 5’ and 3’ flanks remain in close proximity (*arrowhead*). Projections of confocal stacks through 0.5 – 1 µm; scale bars, a, *5 µm*; b-d, *2 µm*. **e**, 25 Mb Hi-C contact map and a 1 Mb zoom view of five genes for cell types, in which respective gene is expressed (*on*) one silent (*off*). TSS and TTS of the genes are marked with light blue lines. **f**, *Cis/trans* ratio profiles, i.e. the total number of Hi-C contacts of a locus with loci on the same chromosome divided by the total number of contacts with other chromosomes, calculated for the gene of interest (*red*) and compared to 9-10 other long genes with low expression (*gray*, see Supplementary Table2 for the lists of genes). For comparability of cis/trans ratio profiles, the x-coordinates are rescaled such that TSS and TTS of all genes are aligned (*shaded areas*). To highlight potential dips localized in the gene body against longer range variations, *cis/trans* ratio profiles are normalized to unity in the region outside the gene body. The *cis/trans* ratio for the *Tg* gene shows a pronounced dip in thyroid; *Ttn* in myotubes and *Myh11* in bladder show a moderate dip; profiles for the less expressed *Cald1* and *Neb* are not significantly different from those in not expressing cells or other long lowly expressed genes.

To further study the arrangement of TLs within their harboring chromosomes we employed Hi-C analysis. We reasoned that if an expressed gene loops out of its chromosome, its *cis*-contacts are diminished compared to the *cis*-contacts of the same gene in a silent state, i.e. the *cis*-to-*trans* contact frequency ratio (further referred to as *cis/trans* ratio) is expected to be low. We performed Hi-C analysis of mouse thyroid, bladder, cultured myoblasts and myoblast-derived differentiated myotubes (Fig.6e). First of all, we observed that *cis/trans* ratios are generally lower in the A than in the B compartment (Extended Data Fig.7), which is in agreement with a previously reported positive correlation between interchromosomal contact probability and transcriptional activity^31-33^, and can be tentatively attributed to the A compartment being enriched in expressed genes looping out from chromosomes.

Secondly, analysis of the *cis/trans* ratios revealed that genes exhibiting the largest TLs, such as *Tg, Ttn* and *Myh11*, display a dip in the *cis/trans* ratio profile, in contrast to the same genes in a silent state or other long but weakly expressed genes. The *Neb* and *Cald1* genes, characterized by lower expression and visibly smaller TLs, have *cis/trans* ratios similar to those of other long and lowly expressed genes (Fig.6f). In agreement with the observation that highly expressed long genes loop out of chromosomes, we found that none of the selected genes significantly altered contacts across them, either in the “*off”*, or in the “*on”* states (Extended Data Fig.8). Thus we concluded that the effect of TL formation is on average localized to the immediate vicinity of a gene and leaves most of the harboring chromosome largely unaffected.

### Transcription Loops are stiff structures

What makes highly expressed genes expand from chromosomes and separate their flanks? Every RNAPII complex is associated with a nascent ribonucleoprotein complex (nRNP) formed by newly synthesized nRNA and numerous proteins involved in RNA processing, quality control, transport, translation, etc.^34^ (Fig.7a1). Therefore, nRNPs containing long nRNAs are voluminous structures^8^ exceeding the size of nucleosomes (10-11 nm^35^) or even that of RNAPII complexes (13-14 nm^36^). Stiffening due to steric repulsion between bulky sidechains is well-known phenomenon in polymer bottlebrushes^37^ and requires sidechains (nascent nRNPs) to be density arranged along the main chain. Since during a transcription burst, RNAPIIs travel along a DNA-template as tightly spaced convoy^27^, we hypothesized that the dense decoration with voluminous nRNPs turn a highly expressed gene into a stiff bottlebrush structure (Fig.7a2; Supplementary Video2).

**Figure 7.**
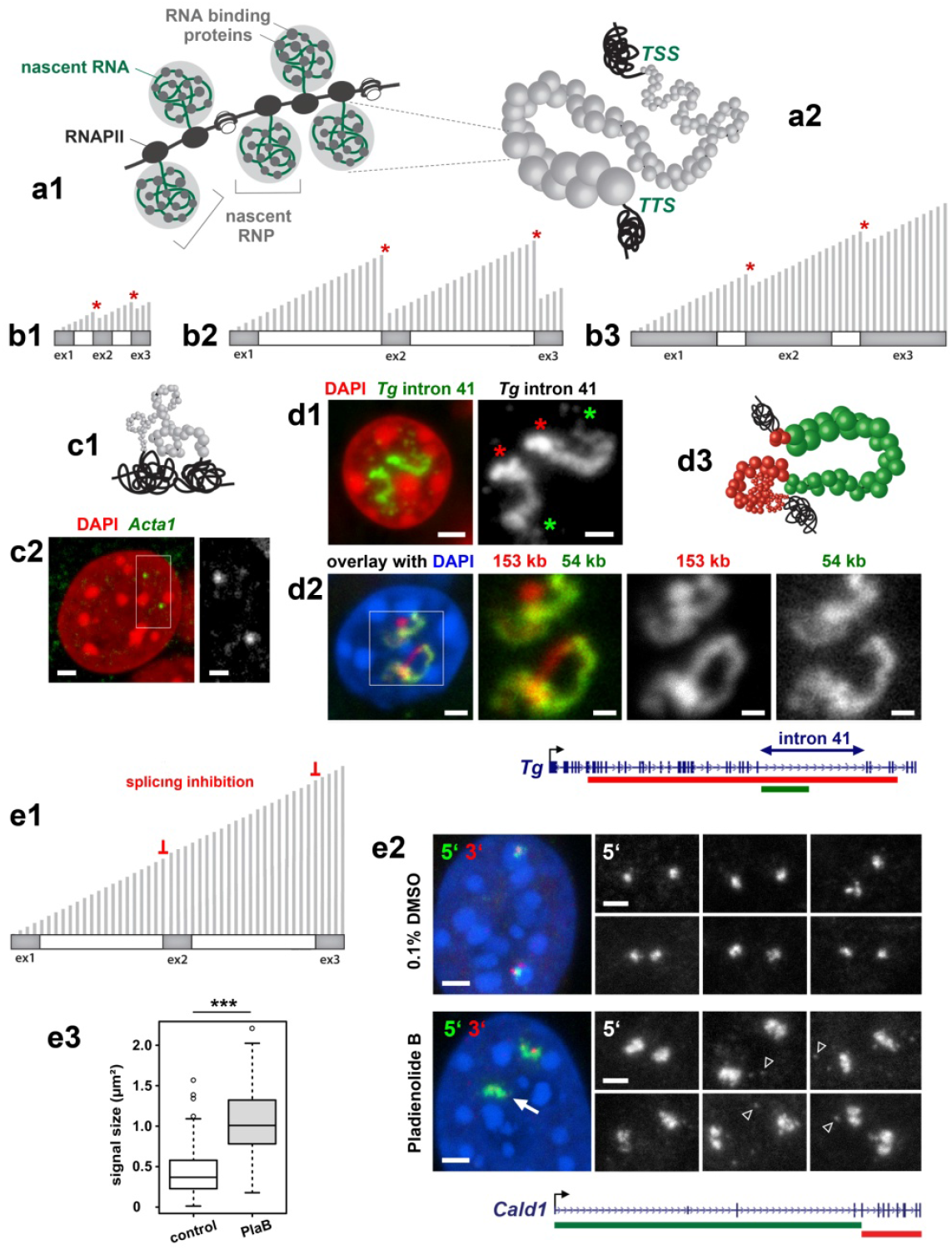
TLs of long highly expressed genes are stiff structures. **a**, Cartoon depicting the formation of nRNPs consisting of nRNAs bound by multiple RNA binding proteins (a1). In long genes, nRNPs decorating a gene body are voluminous. Due to high expression, a gene axis densely decorated by nRNPs loses its flexibility, acquires stiffness and forces the gene to expand (a2). **b**, Schematics showing a short gene (b1), a long gene with long introns (b2) and a long gene with long exons (b3). Exons and introns are shown as *dark-grey* and *white rectangles*, respectively; transcripts are depicted as perpendicular *light-grey lines* of only half of the template length; splicing events are marked with *red asterisks*. **c**, Cartoon showing a short gene decorated by small RNPs, which allows the gene axis to remain flexible and coil (c1). Although the short *Acta1* gene (3 kb) is highly expressed in muscle (4,360 TPM), its RNA-FISH signal remains fairly small (c2). **d**, The *Tg* gene includes relatively long intron 41 (54 kb), which strongly expands and exhibits a gradient of nRNPs from 5’ (*green asterisks*) to 3’ (*red asterisks*) splice sites (d1). Extension of the intron (*green*) is disproportional in comparison to the rest of the gene (*red*, d2) as depicted on the cartoon (d3). **e**, Schematic showing effect of splicing inhibition on length of nRNAs (e1). Comparison of *Cald1* TL size signal in control myoblasts (0.1% DMSO) and myoblasts treated with splicing inhibitor Pladienolide B (10nM). RNA-FISH signals of 5’ and 3’ ends are shown in *green* and *red*, respectively (e2). RGB images are supplemented with representative gray-scale images of the 5’ gene end, which includes long introns. In addition to splicing inhibition, Pladienolide B causes abortion of transcription that is evident from absence of 3’ signal in a large proportion of *Cald1* alleles (*arrow*) and accumulation of nucleoplasmic granules detected mostly by 5’ probe (*arrowheads*). Despite massive abortion of transcription, the size of RNA-FISH signals was increased 2.5-fold indicating larger expansion of the TLs (e3; \*\*\**p-value < 0*.*001, Wilcoxon rank sum test)*. Distributions of used probes in respect to the studied genes are depicted above the image panels. Scale bars: 2 *µm*, in close ups, 1 *µm*.

Short genes produce short nRNAs (Fig.7b1) corresponding to minute nRNPs, which cannot generate sufficient bottlebrush stiffness to prevent gene coiling even when a gene is highly transcribed (Fig.7c). In contrast, long genes produce long nRNAs corresponding to large nRNPs, with large steric repulsion that induces stiffness, forcing gene expansion. Large nRNPs can be formed on genes with either long introns (Fig.7b2), or multiple long exons (Fig.7b3). The first case can be exemplified by the *Tg* gene with 54-kb intron, over which mRNAs grow from ca. 6 to 60 kb and form a signal gradient (Fig.7d1). More importantly, the intron displays a disproportionately larger extension in comparison to the rest of the gene (Fig.7d2,3), in agreement with its anticipated greater stiffness. The second case can be exemplified by the *Ttn* gene, from which ca. 102 kb mRNA is read: the *cis/trans* ratio curve for *Ttn* displays an asymmetrical drop towards the 3’ end (Fig.5f), indicating a stronger exclusion of the 3’ gene end from the harboring locus.

Next, we used splicing inhibition to increase length of nRNAs experimentally, (Fig.7e1), reasoning that this will lead to an increased size of nRNPs and thus to increased stiffness and consequently to a stronger TL expansion. Therefore, we inhibited splicing with Pladienolide B^38^ in myoblasts and performed RNA-FISH with probes for the intron-rich 5’ end and the exon-rich 3’ end of the *Cald1* gene (Fig.7e2). Despite the massive abortion of transcription during drug treatment, we indeed observed a 2.5-fold increase of the signal size corresponding to *Cald1* TL parts formed by the intron-rich region of the gene (Fig.7e3).

### Polymer modeling confirms the stiffening is a possible mechanism of transcription loop formation

We next turned to polymer modeling aiming to understand whether stiffening and lengthening of a highly-transcribed gene can give rise to observed TL. We modeled 50Mb region of a chromosome by a polymer made of 1 kb monomers, each corresponding to ∼5 nucleosomes arranged in a 20 nm globule. We simulated 6 territorial chromosomes confined to a spherical nucleus by initiating them from a mitotic-like conformation and letting them expand (Fig.8a; Extended Data Fig.9a). On each chromosome, we assigned a 100 kb region as the gene of interest (Fig.8a,c) and explored at which simulation parameters we can recapitulate the biological observables for the *Tg* gene, including appearance of TLs and distances between flanks measured by microscopy and *cis/trans* contact ratio along the gene from Hi-C (Fig.8b).

**Figure 8.**
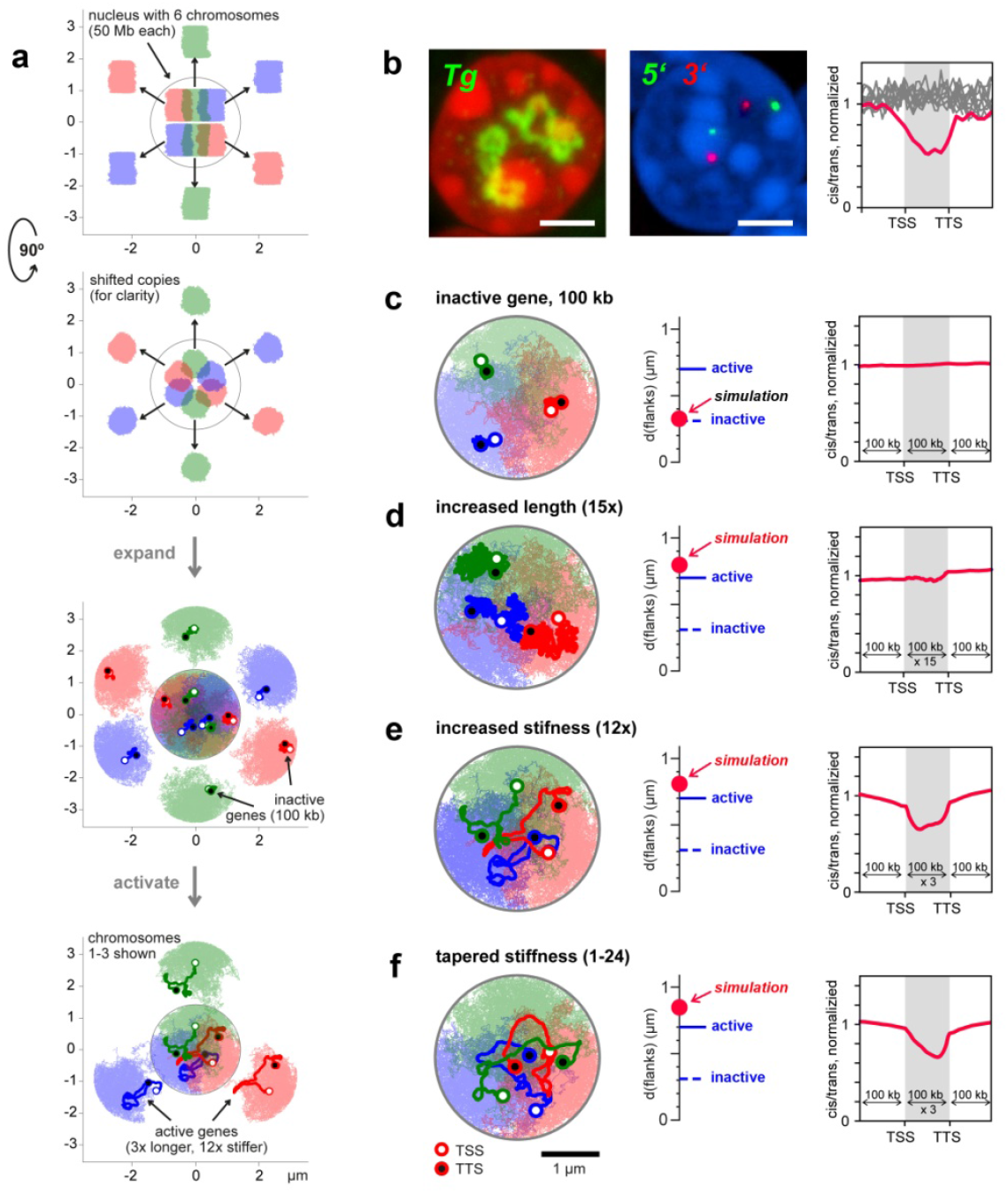
Polymer modeling of transcribed genes. **a**, Polymer simulation setup. Six territorial chromosomes (50 Mb each) in a spherical nucleus were obtained by initiating them in a mitotic-like state (*top and side view*). For clarity, shifted copies are shown outside the nucleus. Then polymers were expanded and for each polymer, a gene of interest of 100 kb was assigned (*depicted with darker color*). For clarity, shifted copies of six chromosomes with simulated genes are shown separately. In the rest of simulations only 3 chromosomes are shown (*bottom*). **b**, Biological variables used to verify TL models: appearance of *Tg* TLs, inter-flank distances from microscopy and *cis/trans* contact ratio calculated from Hi-C maps. **c**, Genes of 100 kb length in the inactive state are compact structures and don’t exhibit a dip in *cis/trans* contact ratio compared to the surrounding fiber. **d**, A mere increase in gene contour length (15-fold) leads to a bigger TL volume but doesn’t exhibit a clearly discernible contour or a dip in *cis/trans* contact ratio. **e**, Increased stiffness (12-fold) together with a moderate increase in contour length (3-fold) shows that TLs expand substantially, exhibit a clear contour, and a dip in *cis/trans* contact ratio. **f**, A gradual increase in stiffness along the gene (in average 12-fold) leads to larger flank separation, more coiling of the 5’ compared to 3’ gene ends and an asymmetric dip in *cis/trans* contact ratio with a steeper slope at the 3’ end.

First, we tried to reproduce TL formation by increasing the gene contour length by 15 fold, reasoning that high expression leads to lengthening *via* complete nucleosome loss (i.e. a 1kb monomer extends to 300nm). However, simulated genes at this condition appeared as rather amorphous structures without discernible contours and thus did not reproduce the visual appearance of TLs. Although median inter-flank distances of simulated genes were comparable to distances measured in thyrocytes (0.88 µm versus 0.75 µm, respectively) the *cis/trans* ratio obtained from simulated Hi-C was not different from the rest of the modeled chromosomes (Fig.8d).

Next, we reasoned that due to dense decoration with nRNPs, a highly transcribed gene has an increased bending stiffness compared to the rest of the chromosome. We increased the contour length by 3-fold to account for partial nucleosome loss and stretching induced by the decoration with nRNPs. Then we altered the stiffness of the simulated genes and found that a 12-fold increase in stiffness reproduces the experimentally observed features of *Tg* TLs (Fig.8e). Moreover, the model successfully recapitulates the dynamic behavior of TL flanks upon gene silencing (Extended Data Fig.9b).

To further account for the gradual increase of nRNA length along the gene, we introduced a tapered stiffness profile along a gene with stiffness gradually increasing from 1 to 24-fold. The added gradient led to a visibly more curled 5’ end and a more stretched 3’ end of the simulated TLs (Fig.8f), recapitulating the morphology of *Tg* TLs observed with microscopy. Furthermore, the stiffness gradient led to asymmetry in the *cis/trans* profile along the gene, in remarkable agreement with experimental Hi-C data (Fig.6f). We point out that in all simulated conditions the dramatic change in visual appearance of the TLs is not accompanied by significant changes in the Hi-C maps with exception to short-range effects. In particular, the formation of TLs does not lead to significant insulation in agreement with experimental data (Extended Data Fig.8).

Importantly, simulations show that a sufficiently dense polymer system (nuclear chromatin) is not confining and allows large-scale relocalization of a long polymer loop (transcribed gene) subject to a sufficiently low force due to stiffening. In conclusion, our simulations serve as a proof of principle that increased stiffness can explain TL formation.

## DISCUSSION

Our study focuses on structure and spatial arrangement of individual transcribed genes in differentiated postmitotic cells and demonstrates that the sole process of transcription can have a profound effect on nuclear architecture. To enable light microscopy observations, we use relatively long highly expressed genes as a model and demonstrate that the selected genes form TLs visually resembling lateral loops of lampbrush^7, 23^ and puffs of polytene chromosomes^8^. The sequential patterns of exon and intron labeling along TLs strongly indicate that RNAPIIs move along the gene axis and carry nRNAs undergoing splicing (Supplementary Video2). This mechanism is drastically different from the popular model of transcription factories with immobilized RNAPIIs and nRNAs extruded in a spot^12, 13^.

We show that transcription dynamically modifies a chromosomal locus harboring an expressed gene: transcription activation causes divergence of gene flanks and gene extension from the chromosome into the nuclear space, whereas transcription silencing causes gene body condensation and convergence of flanks. The fact that TLs are open loops with separated flanks argues against the proclaimed necessity of TSS - TTS association for maintenance of a high level of transcription^39-41^. Moreover, the complicated geometry of highly extended TLs is not compatible with the idea of a perpetual contact between a promoter or an enhancer-promoter complex with a gene body during transcription^42, 43^.

The spreading of TLs over euchromatic nuclear areas by extending away from harboring chromosomes and even breaking them apart, challenges the significance of chromosome territoriality in transcription regulation^30^. A greater number of Hi-C trans-contacts for both euchromatin in general and long highly expressed genes, in particular, indicates that expressed genes can extensively interact with the surrounding euchromatin, suggesting that chromosome territoriality is not an essential functional feature of the interphase nucleus but likely a mere consequence of the last mitosis^5^. Furthermore, observed relocation of a gene by several microns, together with polymer simulations, suggest that the nuclear environment is not confining but allows large-scale chromosomal movements, thus arguing against proposals that interphase chromatin as a gel^44^ or a solid^45^.

We present evidence that highly expressed genes can generate considerable mechanical forces sufficient to re-localize them in a transcription-dependent manner. To explain such a displacement, we hypothesize that TLs are characterized by increased stiffness caused by a dense decoration of the gene axis with progressively growing nRNPs. The hypothesis is supported by differential extension of TL regions with different size of nRNAs and by modeling a transcribed gene as a region of increased bending rigidity. This hypothesis also explains why highly expressed but short genes do not form resolvable TLs: a short gene is lacking long introns or exons and thus decorated by small nRNPs permitting gene-axis coiling (Fig.7). In conclusion, we argue that while formation of microscopically resolvable transcription loops can be specific for long and highly expressed genes (Fig.4), the mechanisms underlying the formation of such loops can be a general aspect of eukaryotic transcription.

## Supporting information

Confocal stacks through nuclei of mouse thyrocytes (counterstained with DAPI, red) after RNA-FISH with genomic probe for the Tg gene (green).

Excel spreadsheet includes distribution of human genes according to their length

Excel spreadsheet includes distribution of mouse genes according to their length

Word table with the list and schematics of probes used for DNA- and RNA-FISH experiments.

Excel spreadsheet containing primer pairs

Information on number of reads in Hi-C experiment and the reproducibility of the replicates.

Cartoon showing how transcription initiation and termination of a highly expressed gene lead to formation and disappearance of a TL.

## FIGURES AND EXTENDED DATA FIGURES

**Extended Data Figure 1.**
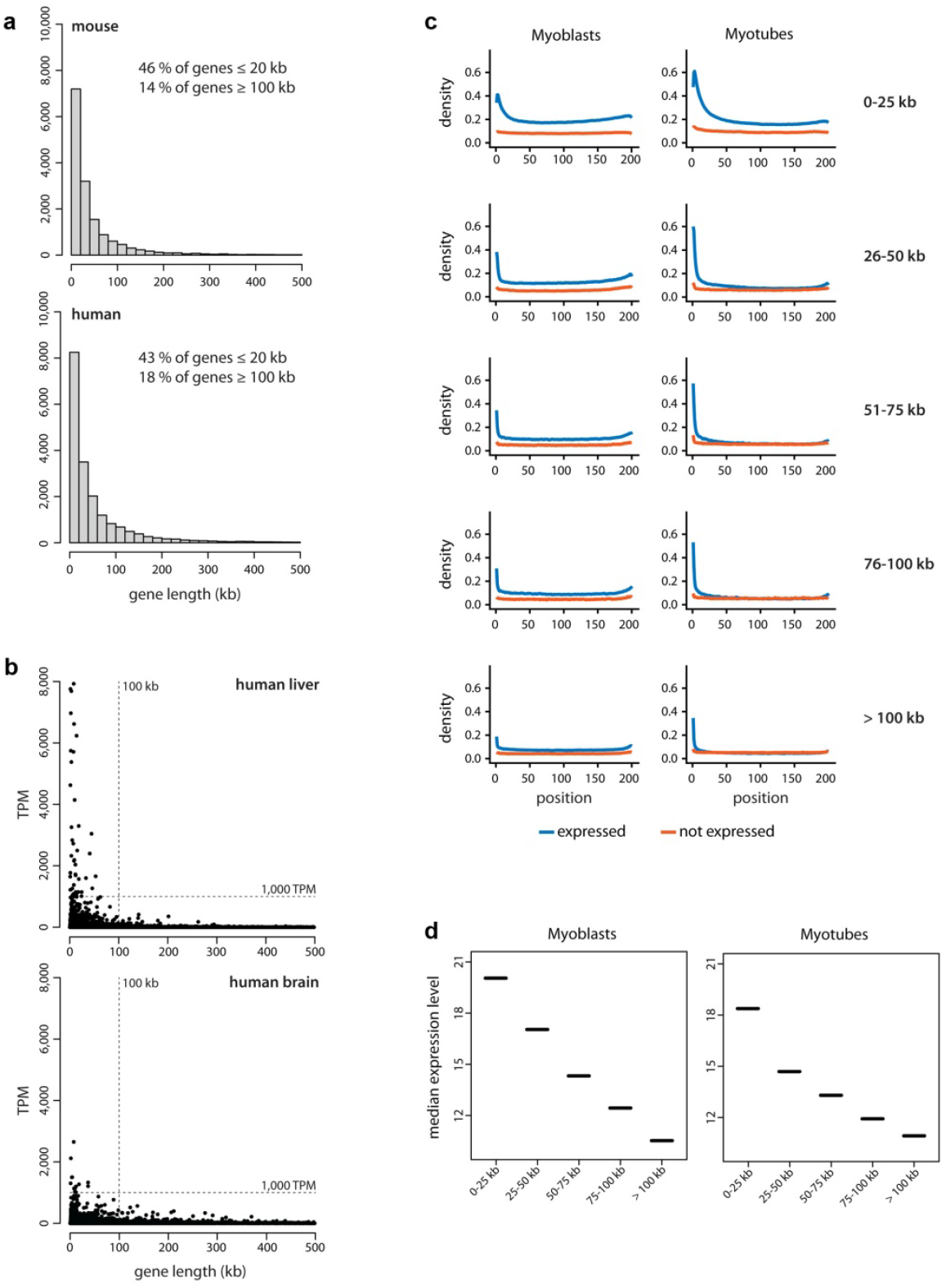
Long genes are rare and expressed at lower levels than short genes. **a**, Analysis of gene length distribution within the human and mouse genomes showed that about 43% and 46% of all protein coding genes, respectively, have a length ≤ 20 kb and only 18% and 14% have a length of 100 kb or above. Bin size: 20 kb. Genes are annotated according to GENCODE. Only genes with a length < 500 kb are shown. **b**, To select suitable genes for visualization with light microscopy, we studied gene expression profiles across 50 human tissues using the publicly available Genotype-Tissue Expression database (GTEx Consortium) and found that long genes are generally not highly expressed. For example, in liver (*top*) and brain (*bottom*) there were no expressed genes with both a length of ≥ 100 kb and with an expression of ≥ 1,000 median TPM. **c**, Comparison of RNAPII occupancy between short and long expressed genes. ChIP-seq with an antibody against the CTD of RNAPII in cultured mouse myoblasts (*left*) and *in vitro* differentiated myotubes (*right*). All genes, expressed (>1 TPM, *blue*) and silent (<1 TPM, *red*), were split into five categories according to their size. RNAPII density (*Y-axis*) is plotted against the respective position within the gene (*X-axis*); each gene is divided into 200 equally sized bins and genes from the same size category are aligned according to the bins. Expressed genes display a higher occupancy with RNAPII compared to non-expressed genes, especially in the TSS region. In the group of expressed genes, the RNAPII occupancy negatively correlates with gene length: the shorter the genes, the higher the RNAPII occupancy. **d**, Analysis of RNA-seq data for myoblasts (*left*) and myotubes (*right*). The median expression level is higher in groups containing shorter genes (<25 kb) and generally negatively correlates with gene length.

**Extended Data Figure 2.**
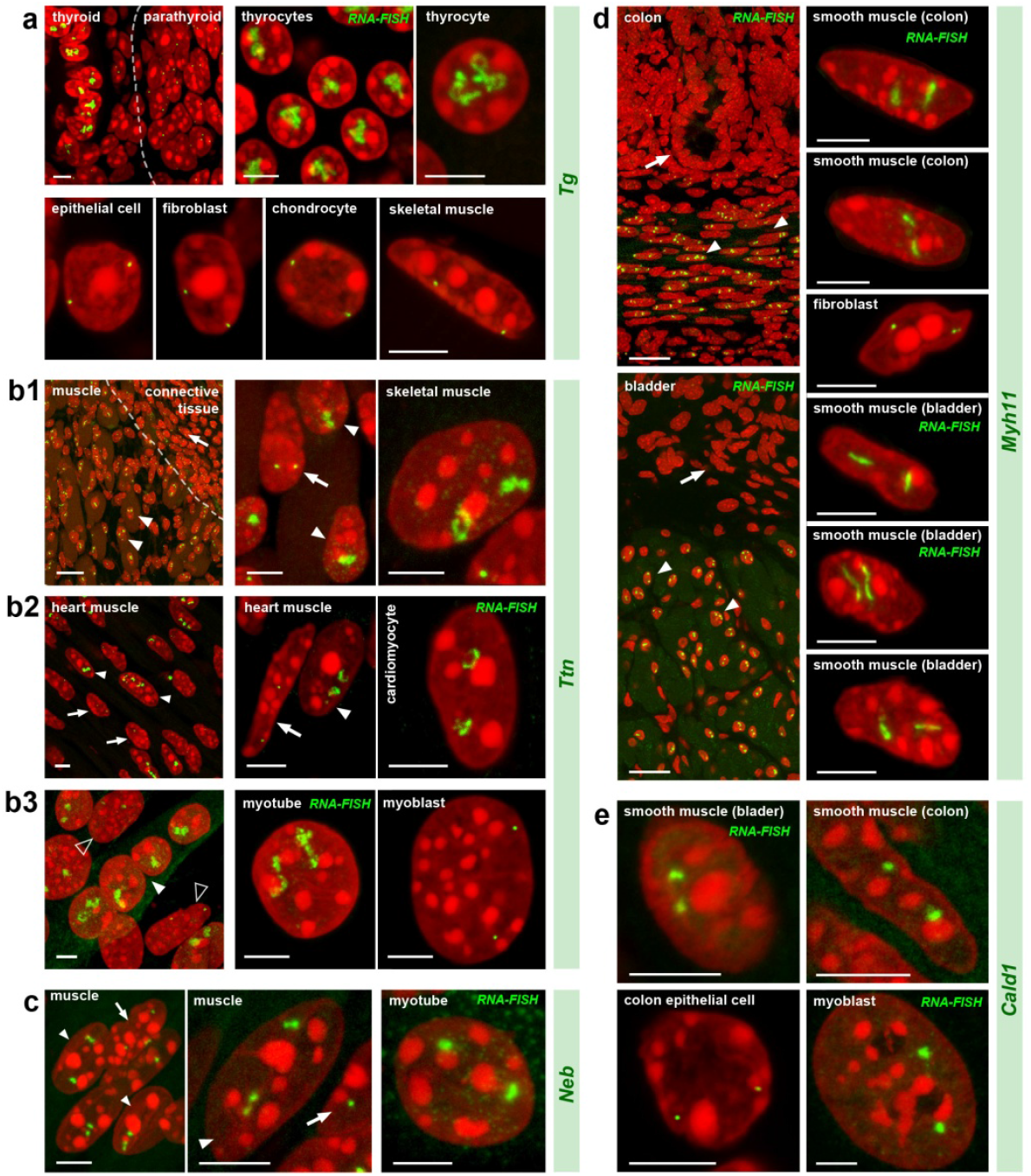
Visualization of the five selected genes in expressing and not expressing cells. **a**, The *Tg* gene is expressed in thyrocytes where both alleles form prominent TLs expanding into the nuclear interior. In neighboring cells with a silent *Tg* gene - parathyroid gland cells, tracheal chondrocytes, epithelial cells, fibroblasts and muscles - *Tg* is highly condensed and sequestered to the nuclear periphery. **b**, The *Ttn* gene is expressed in skeletal muscle (b1), heart muscle (b2) and myotubes differentiated from Pmi28 myoblasts *in vitro* (b3). Note that only muscle nuclei (solid arrowheads) exhibit TLs. In muscle fibroblasts (arrows) or undifferentiated cultured myoblasts (empty arrowheads), *Ttn* is condensed at the nuclear periphery. **c**, The *Neb* gene is expressed in skeletal muscles and cultured myotubes, although to a lesser degree than *Ttn*. Accordingly, it forms smaller TLs. Arrowheads indicate muscle nuclei; arrows indicate fibroblast nuclei with silent *Neb*. **d, e**, The *Myh11* (d) and *Cald1* (e) genes are expressed in smooth muscles of colon and bladder where they form TLs. Note that after RNA-FISH, only smooth muscles (*arrowheads*) but not the neighboring epithelial cells (*arrows*) exhibit TLs. In addition, *Cald1* is expressed in cultured myoblasts and forms small TLs in these cells. Most of the panels display FISH with simultaneous detection of DNA and RNA; panels with only RNA-FISH are marked. All images are *projections of 1-3 µm* confocal stacks. Scale bars for overviews of skeletal muscle, colon and bladder, *50 µm*; for the rest of the panels, *5 µm*.

**Extended Data Figure 3.**
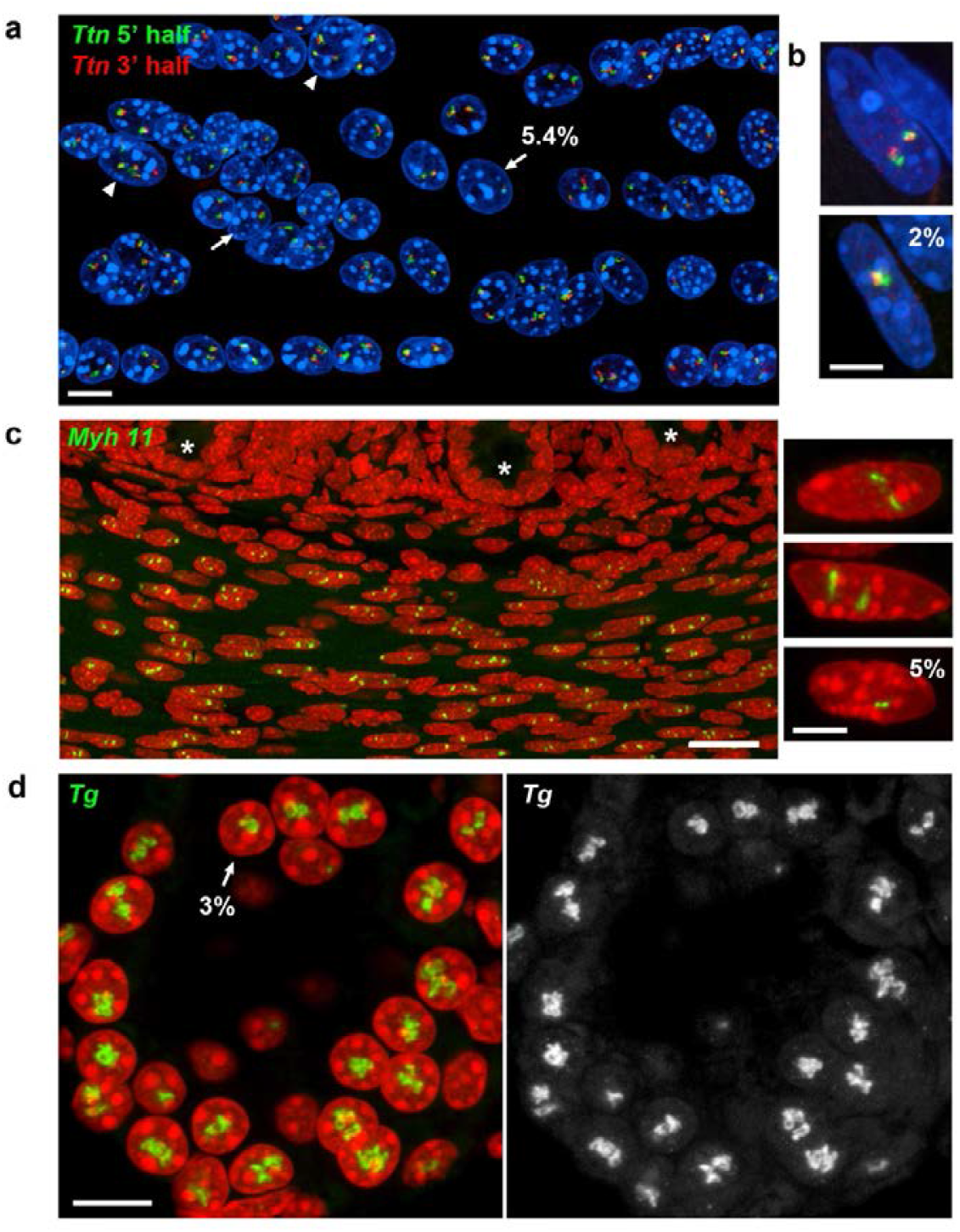
Highly expressed genes are expressed biallelically. Examples of biallelic and monoallelic expression of the *Ttn* gene in cultured myotubes (**a**) and skeletal muscle (**b**), the *Myh11* gene in smooth muscles of intestine (**c**) and the *Tg* gene in thyroid gland (**d**). RNA-FISH with genomic probes. Proportion of nuclei with monoallelic expression is indicated. *Arrows* point at the nuclei with one transcribed allele; *arrowheads* on (a) point at tetraploid myotube nuclei (4 signals); *asterisks* in (c) mark intestinal crypts; note that epithelial cells and fibroblasts do not exhibit *Myh11* RNA-FISH signal. Projections of confocal stacks through 2-4 *µm*. Scale bars: a,d, *10 µm*; b,c (*insertion*), *5 µm*; c (*overview*), *50 µm*.

**Extended Data Figure 4.**
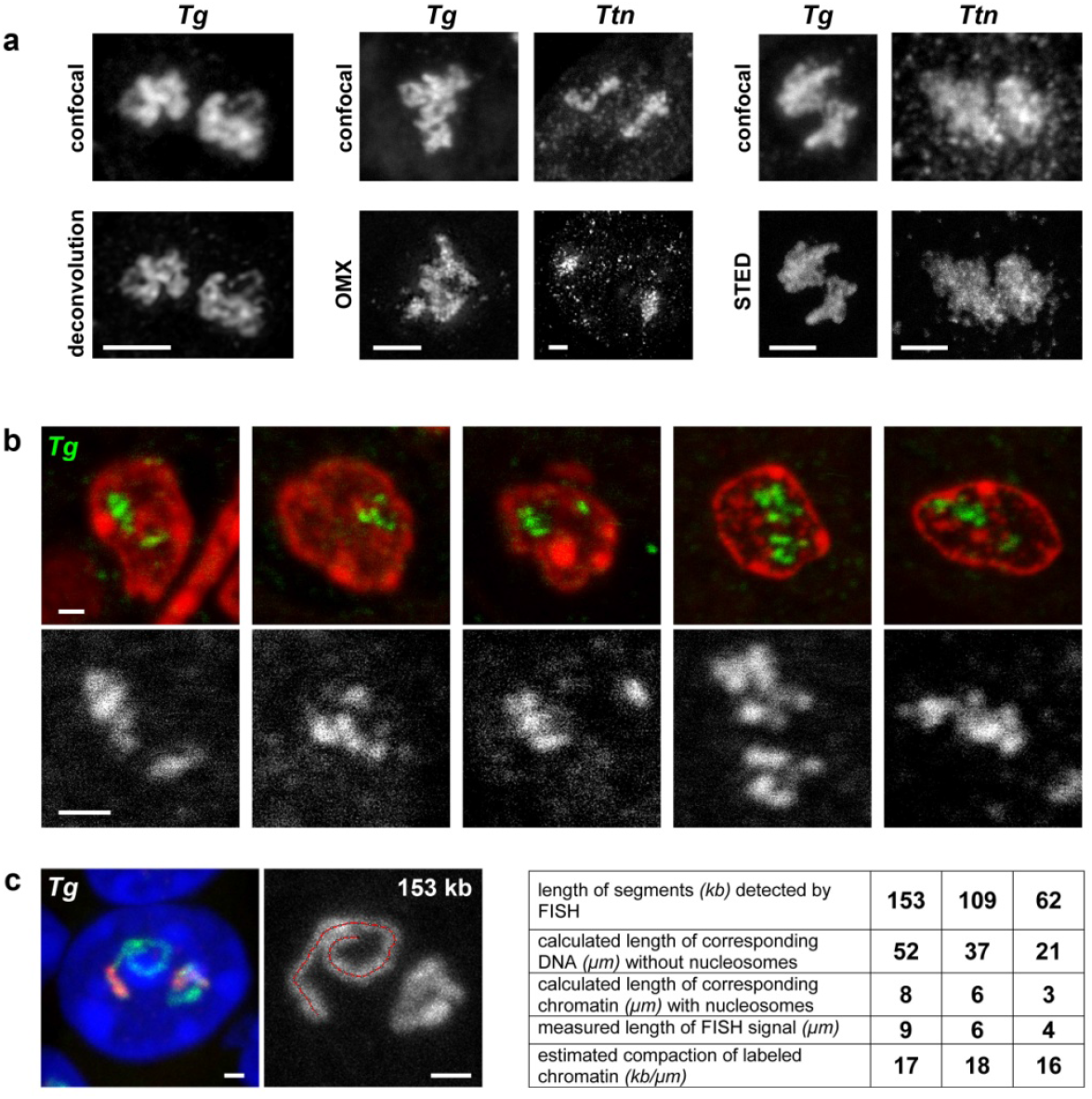
Structure and compaction of TLs. **a**, The internal structure of *Tg* and *Ttn* TLs is not resolvable after deconvolution (*left*) and high resolution microscopy (*mid, right*). **b**, Coiling and folding of TLs is detectable on 50-70 nm thin resin sections. The upper panel shows thin sections through nuclei of thyrocytes with *Tg* TLs (*green*) detected by RNA-FISH. The lower panel shows 2-fold close ups of the corresponding *Tg* TLs images, exhibiting curling and twisting. Images are single optical sections. **c**, To assess the compaction level of TLs, the contour length of three *Tg* TL regions was measured on projections after RNA-FISH. The track of the *Segmented Line tool* in ImageJ, used for measurements, is shown on the *right panel. Tg* regions of 153 kb, 109 kb and 62 kb had a similar compaction level and measured 9 µm, 6 µm and 4 µm, respectively. These values correspond to a nucleosomal structure of chromatin (*table*). However, since *Tg* TLs display internal structures and since the measurements were performed on maximum intensity projections, the compaction level of *Tg* TLs is rather overestimated. Scale bars: a, *2 µm*; b,c, *1 µm*.

**Extended Data Figure 5.**
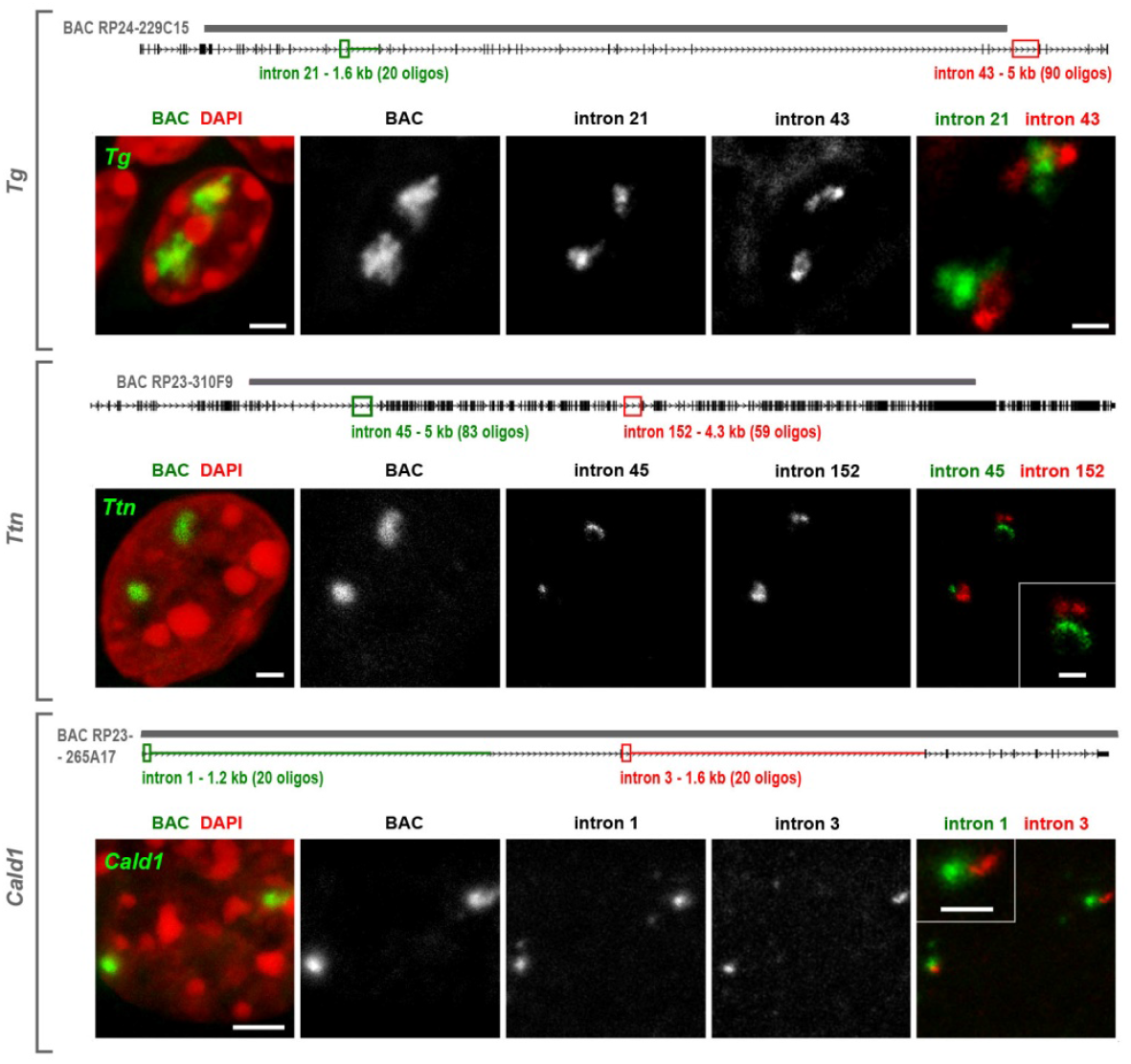
TLs manifest co-transcriptional splicing. Two sequentially positioned introns were labeled with oligoprobes encompassing 1.2 - 5 kb. Schematics above the panels depict distribution of oligoprobes (*green and red rectangles*), labeled introns (*green and red lines*) and positions of BAC probes (*grey lines above genes*). Intron probes label TLs only partially and sequentially. Since the 5’ and 3’ intron signals do not overlap, the 5’ introns are spliced before 3’ introns are read. For instance, in *Cald1*, introns 1 and 3 are separated with intron 2, suggesting that the “green” intron 1 is spliced out before polymerases reach the “red” intron 3, most likely after RNAPII runs over the 3’ splice-site of the first intron. Projections of confocal sections through 2 – 3,5 *µm*. Scale bars: *2 µm*, in close-ups, *1 µm*.

**Extended Data Figure 6.**
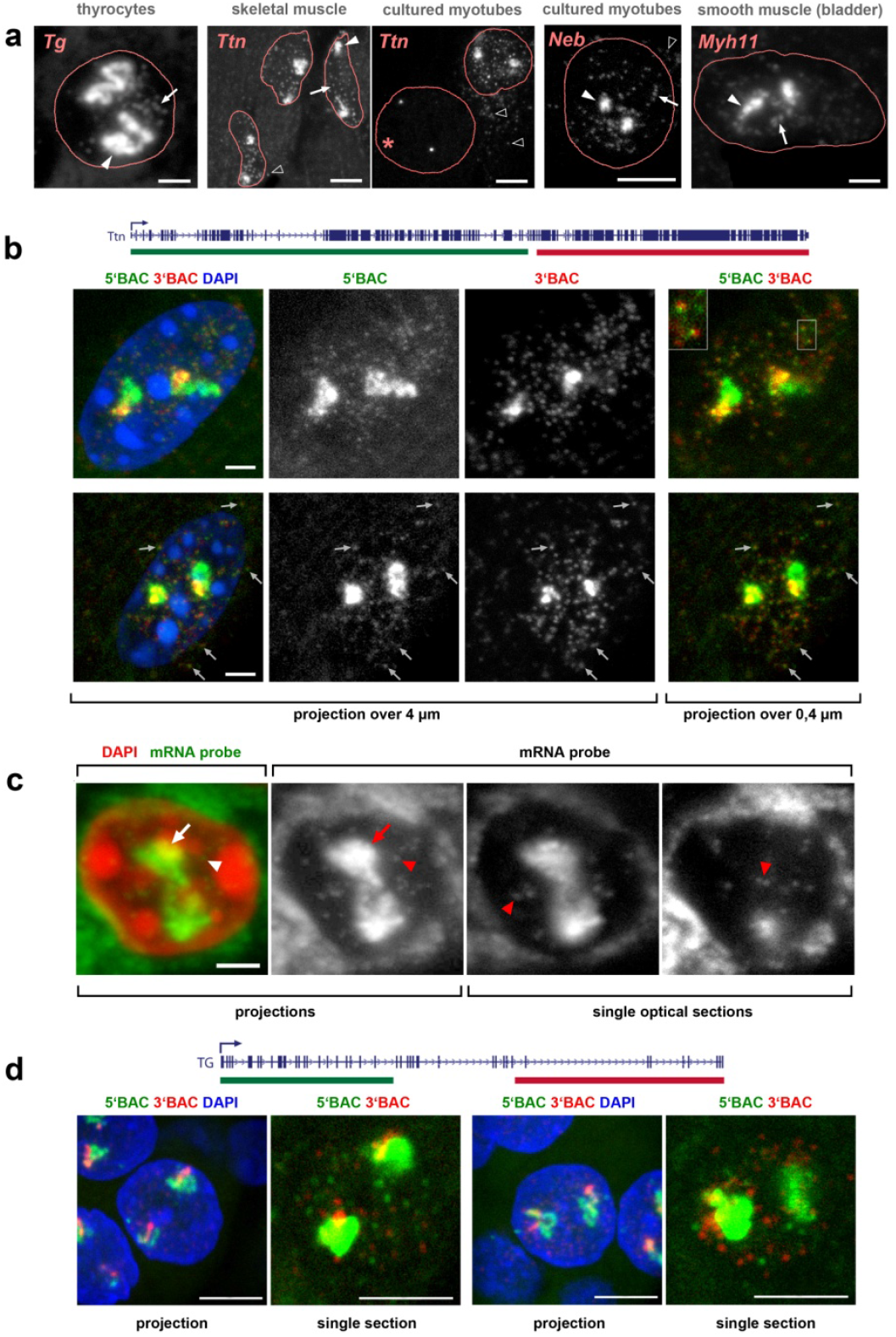
Nucleoplasmic granules in cells with highly expressed long genes. **a**, RNA-FISH reveals numerous nucleoplasmic granules (*arrows*) surrounding TLs (*arrowheads*) in thyrocytes and muscle cells after hybridization with genomic probes. For clarity, only RNA signals are shown within nuclear outlines. Empty arrowheads point at similar granules in myotube cytoplasm. Asterisk marks nucleus of a myoblast not expressing *Ttn*. **b**, Skeletal muscle nuclei after RNA-FISH with 5’ and 3’ *Ttn* BAC probes. The majority of nucleoplasmic granules (81%) are labeled with both probes suggesting that they represent mRNAs. This suggestion is supported by the fact that double-labeled granules are found also in cytoplasm (*arrows on the lower panel*). Moreover, the 5’ and 3’ signals do not entirely overlap (*insertion*), indicating that the two differentially labeled halves of 102 kb long mRNA can be spatially resolved. **c**, In difference to *Ttn*, other long genes have small mRNAs that are not detectable with genomic probes but require exon-specific oligoprobes. The oligoprobe for all 48 *Tg* exons hybridizes to nRNAs decorating TLs (*arrows*) and labels multiple nucleoplasmic granules (*arrowheads*). **d**, Majority of other thyrocyte nucleoplasmic granules are labeled with either 5’ (*green*) or 3’ (*red*) genomic probes with only 10% of granules being double-labeled. Such differential labeling of nucleoplasmic granules, exemplified here for thyrocytes, is characteristic for other studied genes and strongly suggests that these granules represent accumulations of excised introns. Distributions of used probes in respect to the studied genes are depicted above the image panels. Scale bars: a, *2 µm* for *Tg* and *Myh11, 5 µm* for *Ttn* and *Neb*; b, c, *2 µm*; d, *5 µm*

**Extended Data Figure 7.**
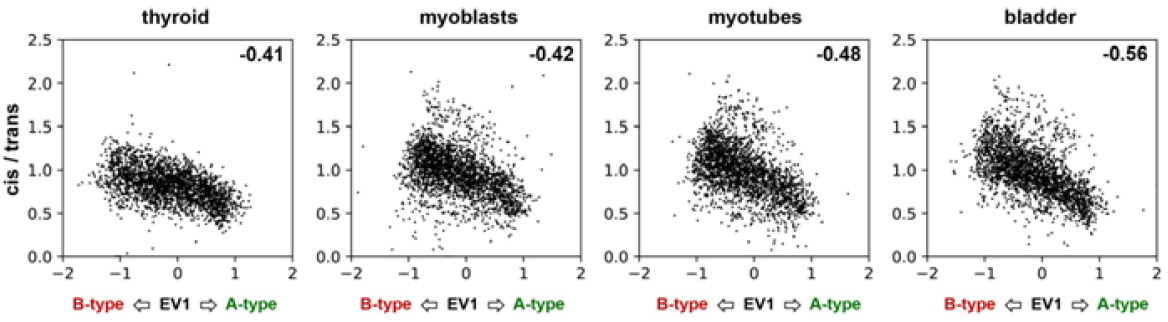
Low *cis/trans* ratios within A compartments in studied cells. *Cis/trans* ratios are lower in A than in B compartments for all four studied cell types. The scatter plots are computed from the genome wide compartment profile and the *cis/trans* ratio profile, both at a bin size of 1,024 kb. The Pearson correlation coefficients are indicated in the upper right corners of the scatter plots.

**Extended Data Figure 8.**
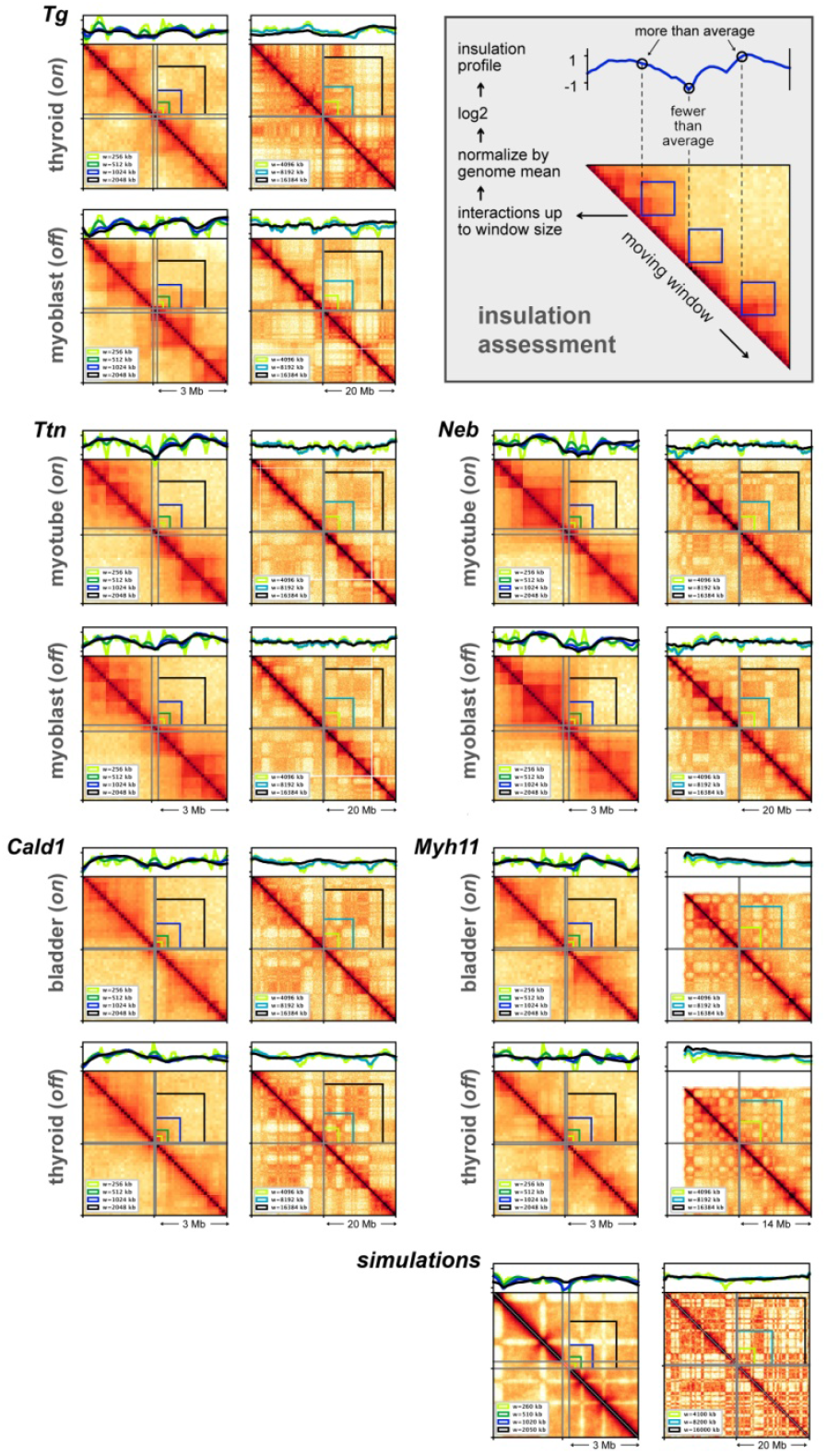
TLs do not cause insulation at different length scales. Insulation^46^ assesses Hi-C contacts spanning across a given locus up to a maximal distance *w* (*top right insert*). Contacts in a square window of size *w* were aggregated and the square was slid along the Hi-C diagonal. The score was normalized by its genome wide mean. Profiles show log2 of the score, such that a locus with profile value −1 has a two-fold reduced number of contacts spanning the locus up to distance *w* compared to the genome wide mean. Insulation scores are computed with the *cooltools* package (https://github.com/mirnylab/cooltools). We computed insulation profiles for Hi-C maps with bin size of 128 kb for various window sizes from 256 kb up to ≈16 Mb. For every analyzed gene, the *left* and *right columns* show a 3 and 20 Mb Hi-C map with insulation profiles for different window sizes; the *top* and *bottom panels* show insulation profiles in expressing (*on*) and not-expressing (*off*) cells, respectively. The analysis shows little correlation between insulation and the formation of TLs: insulation profiles at the gene loci do not differ much between cell types with the gene *on* or *off*. For example, the *Tg* gene shows a moderate dip at scales up to ≈ 1Mb in both thyroids (*on*) and myoblasts (*off*), and no dip in either cell type on the larger scale. Analysis of simulated TLs (see Figure 8 and Extended data Figure 9) confirmed that TL formation does not cause large scale insulation (*bottom row*).

**Extended Data Figure 9.**
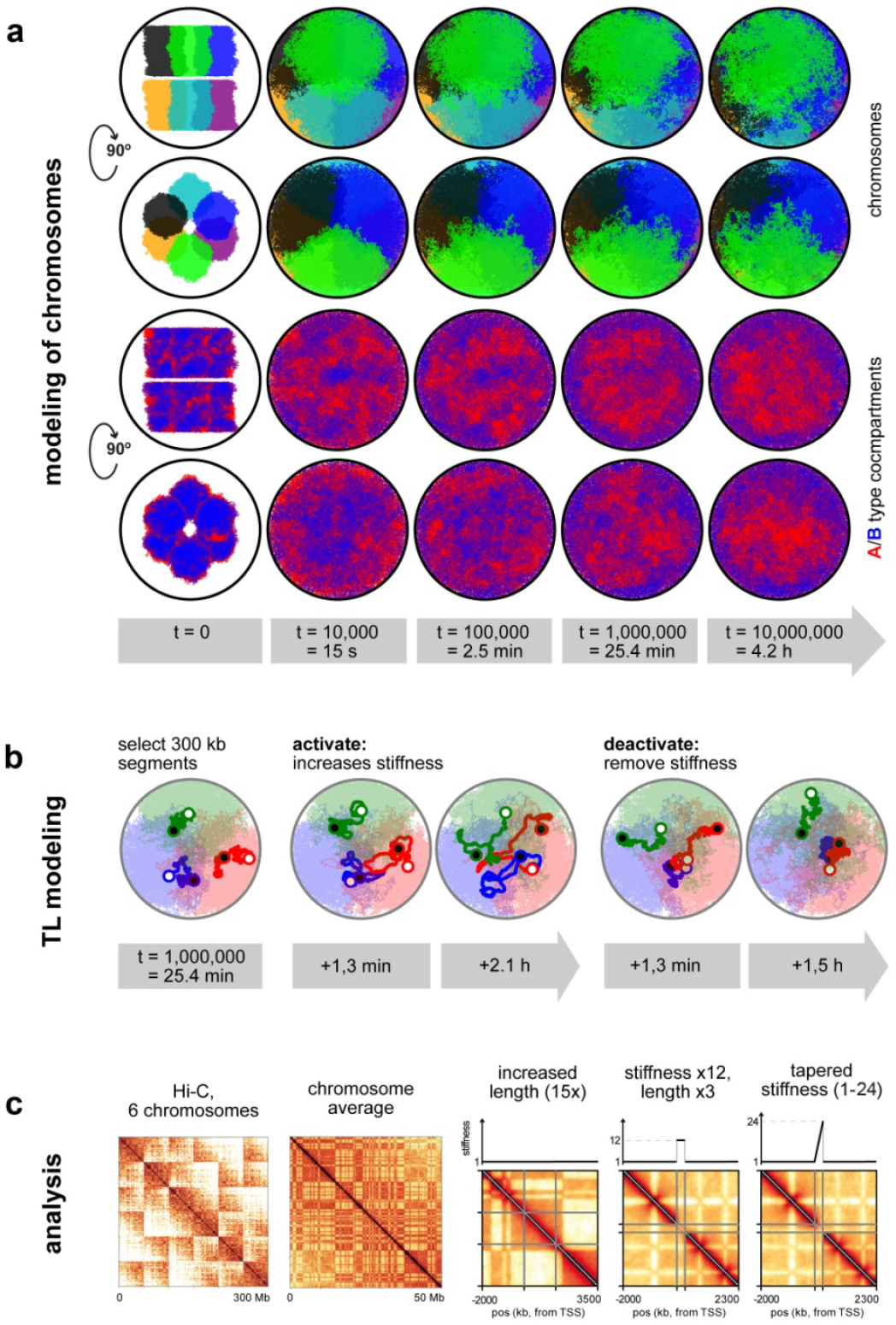
Polymer simulation of chromosomes. **a**, Six chromosomes (50 Mb each) were initiated in a mitotic-like state with unit volume density. Row 1 and 3 show top views, row 2 and 4 show side views. In rows 1 and 2 six chromosomes are differentially colored; in rows 3 and 4 compartmental segments of A and B type chromatin are differentially colored with *red* for A and *blue* for B compartments. The initial expansion is very fast (column 2). However, once the chromosomes fill the nucleus uniformly, subsequent dynamics is very slow and chromosomes retain their territoriality (note that times increase logarithmically). Nevertheless, due to attraction of B-type chromatin to the lamina, a radial structure emerges (rows 3 and 4). **b**, TLs are modeled by choosing a 300 kb segment on each chromosome 25.4 minutes after expansion and increasing the stiffness of the polymer fiber. The genes quickly expand on the order of minutes and were simulated for approximately 1.5 h. The measurements of inter-flank distances and Hi-C maps were performed using configurations sampled from the second half of this time interval. When genes are deactivated by removing the excess stiffness, they collapse back to the inactive state. **c**, Left: Hi-C of all 6 chromosomes shows their territoriality as patches. Second-left: A Hi-C contact map averaged over all 6 chromosomes exhibits the checkerboard pattern typical segregation of A- and B-type chromatin. The three rightmost graphs show zoom views of our genes with stiffness profiles above the maps.

## ACKNOWLEDGEMENTS

We are grateful to David Hörl and Joel Ryan for help with ImageJ plugins and programming. We are thankful to Andreas Maiser and Katharina Brandstetter for help with high resolution microscopy. We acknowledge Jack Bates, Dirk Eik, Peter Becker, Carmo Fonseca, Ada & Don Olins for fruitful and insightful discussions. We thank Dimitar Kralev and Tamako Suzuki for help with animation preparation. This work has been supported by the Deutsche Forschungsgemeinschaft grants (SO1054/1 and SP2202/SO1054/2 to IS, SPP 2202/LE721/17-1 to HL, and SFB1064 to IS and HL) and by National Institutes of Health grants (HG007743 to HL; HG003143 to JD; GM114190 to LM by the Center for 3D Structure and Physics of the Genome of NIH 4DN Consortium, DK107980). JD is an investigator of the Howard Hughes Medical Institute. This paper is dedicated to the memory of Herbert Macgregor who understood loops.

## AUTHOR CONTRIBUTION

IS conceived the project. SLe, SU, YF, KT, IS obtained biological samples. IS, YF, SLe, SU conceived and performed microscopic experiments and image analysis. SLe, CM and SLi performed RNA-seq and ChIP-seq experiments; SB performed RNA-seq and ChIP-seq analyses. YF and EH performed Hi-C experiments; EH, JD, JR and LM performed Hi-C analysis. JR with contribution from LM performed simulations. SLe, SU, JR and IS prepared the figures. IS wrote the manuscript with contributions from SLe, YF, SU, JR, KT, EH, HL, JD and LM.

## COMPETING INTERESTS

The authors declare no competing interests.

## SUPPLEMENTAL INFORMATION

**Supplementary Table1**. Excel spreadsheet includes distribution of human genes according to their length, the list of protein-coding genes and their expression levels in different human tissues, and the list of selected genes with length longer than 100 kb and expression level of ca. 1,000 TPM.

**Supplementary Table2**. Excel spreadsheet includes distribution of mouse genes according to their length, the list of protein-coding genes and their expression levels in two mouse tissues and two cultured cell types, list of five selected genes with length of ca. 100 kb and expression level of ca. 1,000 TPM and list of long lowly expressed genes used for comparison of *cis/trans* contact ratios with selected genes

**Supplementary Table3**. Word table with the list and schematics of probes used for DNA- and RNA-FISH experiments.

**Supplementary Table4**. Excel spreadsheet containing primer pairs used for (1) introduction of protospacer sequences into U6-gRNA-GFP-H2A; (2) generation of oligoprobes for the HD-FISH for *Tg* 41 intron; (3) generation of oligoprobes for SABER-FISH.

**Supplementary Table5**. Information on number of reads in Hi-C experiment and the reproducibility of the replicates.

**Supplementary Video1**. Confocal stacks through nuclei of mouse thyrocytes (counterstained with DAPI, *red*) after RNA-FISH with genomic probe for the *Tg* gene (*green*). Note the volatile shape of the *Tg* TLs and their great expansion into the nuclear interior.

**Supplementary Video2**. Cartoon showing how transcription initiation and termination of a highly expressed gene lead to formation and disappearance of a TL.

The sequence of events: a gene body (*orange thread*) is coiled within a locus (*blue threads*) in a compact structure; upon transcription initiation, RNAPIIs (*dark grey oval structures*) are loading at the gene promoter (*red*) and begin to elongate; during elongation, nRNPs appear and grow in size (*depicted as grey amorphous structures*); during a transcription pause, chromatin of the gene is coiled and forms a sliding knot (*orange thread*), dynamically formed beyond the last RNAPII of the first burst and disentangled by first RNAPII of the second burst; dense decoration with voluminous nRNP rigidifies gene axis and forces its expansion, as well as divergence of gene flanks (*blue threads*); RNAPIIs of the first transcription burst reach the 3’ gene end, release attached nRNPs and dissociate from the gene; the sliding chromatin knot reaches the 3’ gene end; RNAPIIs stop loading at the promoter at the end of the second burst; gene condensation starts 5’-terminally; the gene flanks begin to converge; the gene returns to its coiled compact state.

### METHODS

### GTEx data analysis

RNA-seq data on human tissues was downloaded from the website of the GTEx portal (https://gtexportal.org/home/datasets, version 7, median gene level TPMs). The Genotype-Tissue Expression (GTEx) Project was supported by the <u>Common Fund</u> of the Office of the Director of the National Institutes of Health, and by NCI, NHGRI, NHLBI, NIDA, NIMH, and NINDS. GENCODE gene annotation version v19 (human) was downloaded from https://www.gencodegenes.org/. Using R (https://www.R-project.org/), mitochondrial genes were excluded from the analysis and only protein-coding genes with a known gene status were selected. Gene lengths were calculated from GENCODE annotated start and end positions of each gene. GENCODE annotations were joined with GTEx RNA-seq data based on the genes’ Ensembl gene ID (Supplementary Table 1).

### RNA-seq

Total RNA from cells or tissues was isolated using the NucleoSpin RNA Kit (Macherey-Nagel) according to the manufacturer’s instructions. Digital gene expression libraries for RNA-seq were produced using a modified version of single-cell RNA barcoding sequencing (SCRB-seq) optimized to accommodate bulk cells^47, 48^ After digestion, purification and reverse transcription, the segments were subjected to Illumina sequencing adapters. During the library preparation PCR, 3′ ends were enriched with a custom P5 primer (P5NEXTPT5, Integrated DNA Technologies). Paired-end sequencing of the Nextera XT RNA-seq library was performed using a high output flow cell on an Illumina HiSeq 1500. In the first read, 16 bases were sequenced to obtain the sample-specific barcodes, while the 50 bases in the second read provided the sequence of the cDNA fragment. As multiple libraries were sequenced in parallel on the flow cell, an additional 8 base i7 barcode read was performed.

RNA-seq libraries were processed and mapped to the mouse genome (mm10) using the zUMIs pipeline^49^. UMI count tables were filtered for low counts using HTSFilter^50^. GENCODE gene annotation version vM11 (mouse) was downloaded from https://www.gencodegenes.org/. Using R (https://www.R-project.org/), mitochondrial genes were excluded and only protein-coding genes with a known gene status were selected to ensure proper annotation and used for further evaluation. Gene lengths were calculated from GENCODE annotated start and end positions of each gene (Supplementary Table 2).

### Cell culture

Mouse myoblast cell line Pmi28 was grown in Nutrient Mixture F-10 Ham supplemented with 20% fetal bovine serum (FBS) and 1% Penicillin/Streptomycin at 37°C and 5% CO_2_. For differentiation, cells were seeded at a density of 8 × 10^4^ cells/cm^2^ and incubated in a differentiation medium (high glucose DMEM supplemented with 1% Penicillin/Streptomycin and 2% horse serum) at 37°C and 5% CO_2_. Differentiation medium was replaced every 2 days. A sufficient number of myotubes formed typically after 3-4 days.

### Manipulation of transcription and splicing

Transcription stimulation was performed using a system previously described^28^. Pmi28 mouse myoblasts were transfected with PB-TRE-dCas9-VPR (purchased from Addgene, ID 63800) and PiggyBac transposase (System Biosciences, #PB200PA-1) expression plasmids at a ratio of 3:1 with Lipofectamine 3000 (Invitrogen) according to the manufacturer’s instructions. Clones that stably integrated dCas9-VPR were selected with medium containing 50 µg/ml hygromycin B (Sigma-Aldrich). Single cells were sorted by fluorescence activated cell sorting (BD Biosciences Aria II Sorter) and cultured for one week in presence of hygromycin B. dCas9-VPR expression was induced with 1 µg/ml doxycyclin for 50 h. Individual clones were tested for dCas9-expression by western blot using a Cas9 antibody (Clontech, dilution 1:1000). For gRNA transfection, 3×10^5^ cells were seeded per 6 well on coverslips and transfected with 2.5 µg of a mix of 6 different gRNA plasmids per target gene (Supplementary Table 4) at equimolar amounts with Lipofectamine 3000 according to the manufacturer’s instructions. To monitor transfection efficiency, a plasmid expressing a plasmid expressing GFP-H2A additionally to the respective gRNA (U6-gRNA-GFP-H2A) was generated. CMV-GFP-H2A was inserted into the pEX-A-U6-gRNA expression plasmid via Gibson assembly. pEX-A-U6-gRNA was synthesized at Eurofins MWG Operon according to^51^. Then U6-gRNA-GFP-H2A was added to the gRNA cocktail. dCas9-VPR expression was induced after 12 h by the adding of 1 µg/ml doxycyclin. After 50 h of doxycycline treatment the relative expression level of *Ttn* was assessed by quantitative real time PCR on a LightCycler 480 II (Roche) using LightCycler 480 SYBR Green (Roche) according to the manufacturer’s instructions with the following primers: Ttn fwd (5’-GCCGCGCTAGATTGATGATC-3’) and Ttn rev (5’-TCTCGGCTGTCACAAGAAGCT-3’).

For transcription inhibition in myotubes, differentiation medium was supplemented with 10 µg/ml actinomycin D (Sigma-Aldrich) for 12 or 24 h, or 25 µg/ml α-amanitin (Sigma-Aldrich) for 24 h, or 100 µM 5,6-dichloro-1-β-D-ribofuranosylbenzimidazol (DRB) (Sigma-Aldrich) for 3 or 6 h. Fixation time points of cells under DRB treatment of after DRB removal included 2 min, 25 min, 50 min, 75 min and 3 h.

For splicing inhibition in myoblasts, the culture medium was supplemented with 10 nM of Pladienolide B (Cayman Chemical) for 4 h before fixation.

### Preparation of samples for ChIP-seq and Hi-C

Pmi28 myoblasts were detached using 0.25 % Trypsin/1 mM EDTA for 3 min and pelleted. For native ChIP-seq, the pellet (roughly corresponding to 15 ×10^6^ myoblasts) was directly snap frozen in liquid nitrogen and kept at −80 °C. For Hi-C, cells were resuspended in medium to a final volume of 10 ml. 666 μl of 16% formaldehyde were added (resulting in a final concentration of 1% formaldehyde); the tube was briefly shaken and then rotated for exactly 10 min at 21 °C. To quench the reaction, 593 µl of 2.5 M glycine were added; the tube was briefly shaken and left rotating for 5 min. After quenching, tubes were centrifuged for 5 min at 2,000 g and the supernatant was discarded. Cells were resuspended in 1.5 ml PBS, transferred into 2 ml tubes, centrifuged at 2,000 g for 5 min at RT and the supernatant was discarded. The pellets were snap-frozen in liquid nitrogen and stored at −80 °C.

Myotube samples were collected as follows in order to separate differentiated myotubes from undifferentiated myoblasts: first, myotubes were gently trypsinized using 0.025 % Trypsin/0.1 mM EDTA; when myotubes detached, leaving the majority of myoblasts still attached to the bottom of the dish, the supernatant containing myotubes and a proportion of detached myoblasts was transferred to a fresh 150 mm cell culture dish. The remaining myoblasts were allowed to re-adhere to the dish bottom, whereas myotubes did not re-adhere and thus could be collected with the supernatant. After centrifugation for 4 min at 1,300 g, cells were washed with PBS and centrifuged again. For native ChIP-seq, the pellet (corresponding to 10 ×10^6^ myotubes) was snap frozen in liquid nitrogen and kept at −80 °C. For Hi-C, myotubes were fixed, quenched and frozen in the same way as myoblasts. We estimated that our myotube samples contained 10-20% of myoblasts.

Mouse thyroids for Hi-C were prepared as follows: thyroid glands from three mice were isolated within 5-10 min and stored in medium containing TSH (DMEM/F-12; 20% FCS; Penicillin/Streptomycin; 2 mM L-Glut; 40 μg/ml vitamin C; 1 U/ml TSH) in a cell culture incubator at 37 °C. Glands were then processed one by one. The two lobes of each gland were cleaned from attached neighboring tissues under a binocular and placed back in the incubator in the same medium while the other glands were being processed. Finally, all six lobes were joined in 2 ml of DMEM without FCS, minced with micro-scissors into small pieces (5-6 from each lobe) and transferred to a 2 ml tube using low retention tips. The tube was centrifuged at 2,000 rpm for 5 min at RT and the supernatant was discarded. Tissue was resuspended in 1.5 ml of DMEM without FCS. 100 μl of 16% formaldehyde were added (resulting in a final concentration of 1% formaldehyde); the tube was briefly shaken and then rotated for exactly 10 min at 21 °C. To quench the reaction, 89 μl of 2.5 M glycine were added; the tube was briefly shaken and left rotating for 5 min. Tube was moved to ice and kept there for 0.5 h. After quenching, tubes were centrifuged for 10 min at 6,000 rpm and supernatant was discarded. The pellets were snap-frozen in liquid nitrogen and kept at −80 °C. The same procedure was repeated with three other mice so that each biological replicate consisted of two tubes each containing cells from 6 thyroid lobes. We estimated that thyroid samples contained ca. 60% of thyrocytes.

For smooth muscles, one mouse bladder per sample was used. The tissue was minced and fixed as described for thyroid. We estimated that bladder samples contained 40-50% of smooth muscle.

### Native ChIP-seq

Native ChIP-seq was performed as previously described^52, 53^ with the following alterations: IPs were carried out with S1 mononucleosomes derived from Pmi28 myoblasts or myotubes. Experiments were performed in independent duplicates, each including. Per one IP reaction (500 µl), mononucleosomes from 5×10^6 cells were incubated with 5 µg of antibody (anti-RNA polymerase II CTD repeat YSPTSPS, 8WG16, Abcam ab817) in rotating tubes at 4°C overnight. Then the antibody were post-coupled to magnetic beads (Dynabeads Protein G, Invitrogen) at 4°C for 3 h and eluted IP DNA was Phenol/Chloroform/Isoamylalcohol extracted using MaXtract high density tubes (Qiagen) and ethanol precipitated according to a standard protocol. Illumina sequencing libraries were prepared using the NEBNext Ultra II DNA Library Prep Kit for Illumina (New England Biolabs) and the NEBNext Multiplex Oligos for Illumina (New England Biolabs) according to the manufacturer’s instructions. ChIP-seq libraries were sequenced on an Illumina HiSeq 1500 platform using 50 bp single end nucleotide sequencing.

### Hi-C

3 Hi-C biological replicates were prepared for each of the studied cell types – myoblasts, myotubes, thyrocytes and smooth muscles. Sample sizes were estimated as following: 15 x10^6 of myoblasts, 10 x10^6 of myotube nuclei, 3 x10^6 of thyroid cells (thyroids of 6 mice); number of cells in a bladder was not estimated. Hi-C was performed as described by^54^. Myoblasts and myotubes were split into 3 or 2 subsamples, respectively, each containing 5 x10^6 cells, which were treated as separate samples until the DNA purification step when they were merged into one biological replicate. Chromatin was digested with the restriction enzyme *DpnII* (New England Biolabs) at 37° C overnight. Chromatin sonication and size selection were performed with some modifications to allow retrieval of a higher DNA quantity. Hi-C libraries were sequenced using 50 bp paired-end sequencing on an Illumina HiSeq 4000. For the number of reads obtained in each Hi-C experiment and the reproducibility of the replicates see Supplementary Table 5.

### Hi-C data processing

Fastq files were mapped to the mm10 genome using the distiller pipeline (github.com/mirnylab/distiller-nf). Hi-C interaction data was stored in the cooler format^55^. Cooler: scalable storage for Hi-C data and other genomically labeled arrays. *Bioinformatics*. doi: 10.1093/bioinformatics/btz540). We always applied iterative correction to our data in order to remove biases as much as possible^56^.

Cis/trans contact frequencies were calculated for each bin as the number of cis (same chromosome) contacts divided by the number of trans (different chromosome) contacts of that bin. This is repeated for all bins along the genome, resulting in a genomic track of cis/trans contact frequency ratios.

For a comparison plot of the cis/trans ratio across a number of genes the x-axis for each gene was rescaled so that TSS and TTS align. Interpolation was used to obtain the same number of points in each gene. The cis/trans ratios were normalized to regions outside the genes (to be precise: the region from TSS-3’ gene length to TSS-0.5’ gene length and TTS+0.5’ gene length to TTS+3’ gene length is used to normalize cis/trans profile to unity). This is done to highlight local changes in the cis/trans ratio associated with genes while suppressing longer-range variations.

### Mice and tissue sampling

CD-1 mice were purchased from Charles River Laboratories, housed at the Biocenter, Ludwig Maximilians University of Munich (LMU) and treated according to the standard protocol approved by the Animal Ethics Committee of LMU and The Committee on Animal Health and Care of the local governmental body of the state of Upper Bavaria, Germany. Mice of both sexes were used with age from P14 to adult. Mice were sacrificed by cervical dislocation after IsoFlo (Isofluran, Abbott) narcosis or by CO_2_ euthanasia.

For isolation of the thyroid gland, a vertical anterior neck incision was made, a piece of trachea (ca. 5 mm) with the attached thyroid lobes was excised and placed into PBS. The thyroid tissue was carefully cleaned from connective and fat tissue, muscles and parathyroid glands under a binocular. In order to relax smooth muscles of a bladder, mice were injected with Narcoren (Boehringer Ingelheim, conc. 5 µl/g body weight) before euthanasia and cervical dislocation. The abdominal musculature was carefully removed, the bladder was excised and placed into PBS. For isolation of colon, a vertical incision in the lower abdomen was made and extended horizontally in both directions, the abdominal musculature was carefully removed and a part of the distal colon was excised.

All isolated tissues were washed with PBS and then fixed with 4 % paraformaldehyde (Carl Roth) solution in PBS for 12-20 h. After fixation, mouse tissues were washed with PBS and incubated in increasing concentrations of sucrose - 10 %, 20 %, 30 % - for 1 to 12 h at +4°C. Tissues were embedded in Tissue-Tek O.C.T. compound freezing medium (Sakura) in the desirable orientation. Blocks were stored at –80°C before cutting into 16-20 µm sections using a cryostat (Leica CM3050S). Cryosections were collected on Superfrost Plus slides (Thermo Scientific) and stored at –80°C before use.

For preparation of thin sections, thyroid glands were fixed with mixture of 2 % paraformaldehyde and 0.1 % glutaraldehyde in 300 mOsm cacodylate buffer (75 mM cacodylate, 75 mM NaCl, 2 mM MgCl_2_) for 30 min and then immediately embedded into Lowicryl. Thin sections (50-70 nm) were prepared using Reichert Ultracut.

### FISH probes

#### BAC-derived FISH probes

BAC clones encompassing the desired genomic region (Supplementary Table 3) were selected using the UCSC genome browser (https://genome.ucsc.edu/) and purchased from BACPAC Resources (Oakland children’s hospital) as agar stabs (https://bacpacresources.org/). BACs were purified via standard alkaline lysis or the NucleoBond Xtra Midi Kit (Macherey-Nagel), followed by amplification with the GenomiPhi Kit (GE Healthcare) according to the manufacturer’s instructions. BAC probes were labeled either directly with fluorophores (using fluorophore-dUTPs), or with haptens (biotin-dUTP, digoxigenin-dUTP) by nick translation.

#### Chromosome paints

Whole chromosome paints were a kind gift from Nigel Carter and Johannes Wienberg (Cambridge). Paints were amplified and labeled with Biotin or Digoxigenin via DOP-PCR with a partially degenerate oligonucleotide primer (6MW primer: 5’-CCG ACT CGA GNN NNN NAT GTG G −3’, Eurogentec)^57^.

#### Tg exon probes

Probes to specifically label exons 2-12 and 33-47 of *Tg* were generated by standard polymerase chain reaction (PCR) using a cDNA clone (transOMIC technologies, BC111467) as template. Primer sequences are listed in the Supplementary Table 4. PCR amplified DNA was directly labeled with fluorophores by nick translation.

#### Oligoprobes

Most of oligoprobes were generated using the original SABER-FISH protocol^58^. BED files for the genomic locations of the exons and introns of the target genes were acquired from the UCSC Genome Browser. The designed oligos were ordered as oligo pools from IDT in a scale of 50 pmol/oligo. All used probe oligos are listed in Supplementary Table 4. The oligos were re-mapped if necessary for multi-color imaging and extended to ∼500 nt using the previously described primer exchange reaction^59^. Finally, the probes were purified using PCR clean-up columns (Macherey Nagel).

#### Oligoprobes for Tg intron 41

FISH probe generation was conducted as described previously for HD-FISH^60^. Probe covering the first half of *Tg* intron 41 was amplified from genomic J1 ESC wt DNA. Primers to amplify fragments for FISH probes were designed with the HD-FISH probe generator platform accessible via http://www.hdfish.eu/^60^ and are indicated in Supplementary Table 4. After amplification, DNA was labeled with Alexa594 or Alexa647 using Ulysis Nucleic Acid Labeling Kits (Invitrogen).

### FISH protocols

Labeling of dUTPs, probe labeling by nick-translation and preparation of hybridization mixtures were performed as previously described^57^. A typical precipitation mixture with a BAC derived probe was set-up as follows: 100 µl BAC−fluorophore/hapten, 25 µl Cot-1 DNA (Thermo Fisher), 3 µl salmon sperm DNA (Thermo Fisher), 500 µl prechilled (−20°C) ethanol. A typical precipitation mixture with PCR derived probes was set-up as follows: 10 µl PCR probe−fluorophore, 10 µl salmon sperm DNA (Thermo Fisher), 250 µl prechilled (−20°C) ethanol.

For 3D-FISH, cells were subcultured on coverslips coated with 1mg/ml poly-L-lysine (Sigma-Aldrich). Coverslips with cells were washed twice with pre-warmed (37°C) PBS and fixed with 4 % PFA (Merck)/PBS for 10 min at RT. During the last minute of fixation, a drop of 0.5 % Triton X-100/PBS was added to prevent cell distortion during solution changes. Cells were washed 3x with PBS/0.01 % Triton X-100 for 5 min and then permeabilized with PBS/0.5 % Triton X-100 for 10 min. Further pretreatments included incubation with 20% glycerol, followed by several cycles of freezing / thawing, and incubation with 50% formamide/2xSSC^61, 62^.

For DNA-FISH, cells were treated with 50 µg/ml RNase A at 37°C for 1 h. For DNA-FISH or FISH detecting both DNA and RNA, denaturation of both probe and sample DNA was carried out simultaneously on a hot block at 75°C for 3 min. For RNA-FISH, only the probe DNA, but not the sample DNA was denatured. Hybridizations were typically carried out in water-bath at 37°C for 2 days. Immuno-FISH was performed as previously described^61^. Briefly, after immunostaining, cells were post-fixed with 2% PFA/PBS for 10 min, incubated in 20% glycerol for 1h and further treated according to the protocol for 3D-FISH.

RNA-FISH with oligoprobes (SABER-FISH) was performed by mounting SABER-probe containing hybridization mixture on fixed cells and incubating O/N at 43°C. The samples were washed twice (x20 min) with 2xSSC at 37°C, 5 min with 0.1xSSC at 60°C, and rinsed with PBS at RT. Hybridized SABER probes were detected by incubating with 1 μM fluorescently labeled detection oligonucleotides in PBS for 1h at 37°C followed by washing with PBS for 10 min.

FISH on cryosections was performed as previously described^63^. Briefly, after 30 min of drying up at RT, slides were heated in sodium citrate buffer at 80°C for antigen retrieval and then incubated in 50 % formamide/2xSSC for minimum 30 min before probe loading. DNA- and RNA-FISH on sections were performed as described for cells. Simultaneous denaturation of probes and sections was carried out on a hot block at 80°C for 3 min. For combined RNA-FISH and immunostaining, the cryosections were first hybridized with probes, then permeabilized with 0.1 % Triton X-100/PBS and incubated with both primary and secondary antibodies as described previously^64-66^.

In all experiments, DNA was counterstained with 2 µg/ml DAPI in PBS for either 10 min (cells), or for 30 min (sections). Coverslips with cells or cryosections were mounted in antifade (Vectashield, Vector) and sealed with nail polish before microscopy.

### Microscopy

Confocal image stacks were acquired using a TCS SP5 confocal microscope (Leica) using a Plan Apo 63/1.4 NA oil immersion objective and the Leica Application Suite Advanced Fluorescence (LAS AF) Software (Leica). Z step size was adjusted to an axial chromatic shift and typically was either 200 nm or 300 nm. XY pixel size varied from 20 to 60 nm. Used laser lines included 405 nm of diode laser (DAPI), 488 nm of Argon laser (FITC, Alexa 488, GFP), 561 nm of DPSS laser (Cy3, TAMRA, Alexa 555, ATTO 565), 594 nm of HeNe laser (Texas Red, Alexa 594, ATTO 590) and 633 nm of HeNe laser (Cy5, ATTO 633, Alexa 647). Obtained confocal stacks were processed using *ImageJ* (https://imagej.nih.gov/ij/). Axial chromatic shift correction, as well as building single grey-scale stacks, RGB-stacks, RGB-montages and RGB maximum intensity projections was performed using ImageJ plugin StackGroom^67^.

Amount of acquired nuclei varied between cell types: *Tg* TLs were visualized in >3,000 thyrocytes; *Ttn* TLs – in >1,000 cultured myotube nuclei, ca. 500 skeletal muscle nuclei and >50 cardyomyocytes; *Neb* TLs – in 80 cultured myotube nuclei and 50 skeletal muscle nuclei; *Myh11* TLs – in >500 smooth muscles of colon and in >300 smooth muscles of bladder; *Cald1* TLs – in 50 smooth muscles of colon, in 90 smooth muscles of bladder and in >500 cultured myoblasts.

For deconvolution of confocal stacks, a Lightning module of SP8 confocal microscope was used. High resolution microscopy of transcription loops after RNA-FISH with BAC-probes was performed using either OMX (DeltaVision OMX V3 3D-SIM microscope, Applied Precision Imaging, GE Healthcare) or with 3D STED (Abberior Instruments). For OMX, nuclei were counterstained with DAPI, probes labeled with Texas Red and samples were mounted in Vectashield. For STED, nuclei were counterstained with SiR DNA, probes were labeled with Alexa594 and samples were mounted into Mowiol. Alexa594 and SiR-DNA were excited with pulsed excitation lasers at 594 or 640 nm, respectively, and depleted with pulsed depletion laser at 775 nm.

### Image analysis

Inter-flank distances were measured using in-house semi-automated program. The signal spots in both channels were identified by detecting local minima in a Laplacian-of-Gaussian (LoG)-filtered image (with the expected size/sigmas set to enhance spots at the diffraction limit). A pairing of spots from both channels with minimal total distance was calculated using linear assignment and coordinates of both partners saved for further analysis. In both the detection and matching steps, results were visualized immediately, with the option to manually curate them and remove erroneous detection of pairings. The spot pair detection was implemented in Python in the form of a Jupyter notebook. The code is available at https://github.com/hoerldavid/fish_analysis.

Contour length of the *Tg* TL regions were measuredon maximum intensity projections of stacks after RNA-FISH. Loop contours were manually traced using the *Segmented Line tool* in ImageJ. For comparison of RNA-FISH signal size of *Cald1* TLs in control cells and after splicing inhibition, maximum intensity projections of stacks after RNA-FISH were prepared and filtered (Gaussian = 1). The signal areas of individual *Cald1* alleles were measured in *Fiji* after applying *Intermodes* threshold.

### Transcription loop modeling

Chromosomes were modeled as polymers with each monomer representing 1 kb or roughly 5 nucleosomes. To fix the three-dimensional (3D) size of a monomer, literature values for the compaction ratio of the chromatin fiber, i.e. the number of bp per nm of contour length were used. Literature values vary greatly and are in the range c = 25 to 150 bp/nm^68-70^. As such the c = 50 bp/nm was used here, which fixed the 3D size 1kb monomers to 20 nm.

For the Kuhn length lK (typical bending radius of the fiber), again, literature values vary greatly and are in the range of lK= 60 to 400 nm (refs as above). Further estimations of the Kuhn length come from our own measurements of spatial distances of the 5’ and 3’ flanks of our genes in the inactive state (Fig.3). Those measurements suggest somewhat smaller Kuhn lengths between 25 and 80 nm. For our polymer model here, we chose lK= 56 nm or 2.8 kb.

Finally, the volume density of chromatin in the cell nucleus, was fixed to 20%. A proxy can be obtained from the genome size (≈ 3 Gb for a single copy, ≈ 6 Gb per nucleus) and the size of the nucleus. With a diameter of 8 µm the volume density is 180 kb in a cube of size (20 nm)^3^. A cryo-EM measurement^71^ suggests volume densities around 30%.

In order to interpret simulations, the simulation time scale is converted to physical time. This conversion is done by comparing mean square displacements of simulation monomers with experimental data on tracked loci in yeast^72^ and mammals^73^ as described^74^. A conversion factor of 656 ps in simulations corresponding to 1s real time was obtained.

Polymer simulations were performed using a Mirny lab written wrapper (available at https://github.com/mirnylab/openmm-polymer-legacy) around the open source GPU-assisted molecular dynamics package Openmm (1,2). Polymers are represented as a chain of monomers with harmonic bonds, a repulsive excluded volume potential, and an additional small attraction for the interaction of two monomers of type B. Furthermore, the association of B-type chromatin with the nuclear periphery was modeled by a weak attraction of B-type monomers with the sphere boundary representing the nuclear envelope.

Six chromosomes of 50 Mb each (corresponding to 50,000 monomers each) were simulated and placed inside a sphere with diameter of 2.84 µm (corresponding to 42 monomer diameters). Territorial chromosomes were generated by initiating them in a mitotic-like conformation and letting them to expand. In a dense environment, polymer dynamics is exceedingly slow, therefore, chromosomes mix only moderately and retain their territoriality over all simulated times, corresponding to at least hours, or days of real time. The above expansion from mitosis was repeated 10 times for 25 min and 24 s (1,000,000 simulation time units) each. These chromosome conformations served as initial conditions for our simulations of long highly transcribed genes.

To simulate the effect of a high expression of a long gene, a 100 kb region on each simulated chromosome was assigned to the gene of interest. In these regions the bending stiffness of the fiber was increased. The “genes” were positioned in a region of A-type chromatin. Unless otherwise indicated, simulations of highly transcribed genes were run for 127 min of real time (5,000,000 simulation time units). In each run, 101 conformations were saved at equidistant time points.

To obtain Hi-C maps from simulated data polymer conformations were coarse grained by a factor of 10 (i.e. only every 10th monomer is considered) in order to reduce the size of the computed Hi-C matrix. A cutoff radius was then designed mimicking the crosslinking radius in an actual Hi-C experiment. The cutoff radius was 10 monomer diameters. Hi-C maps were computed from conformations in the second half of each run. *Cis/trans* contact frequency ratio tracks were computed as for experimental Hi-C matrices. The *cis/trans* ratio tracks of all six simulated chromosomes were aggregated to improve statistics.

## Notes

### Competing Interest Statement

The authors have declared no competing interest.

