## Supplementary material for "Spatial organization of transcribed eukaryotic genes": Word table with the list and schematics of probes used for DNA- and RNA-FISH experiments.

**Supplementary Information Table S3. BAC clones used for DNA- and RNA-FISH in this study**

| ***Tg*** | |
| --- | --- |
| Gene body | RP24-229C15 |
| Gene halves | RP23-193A18, RP23-266I10, RP24-179P7, RP23-255H1 |
| Gene flanks | RP24-171I15, RP24-312F21 |
| 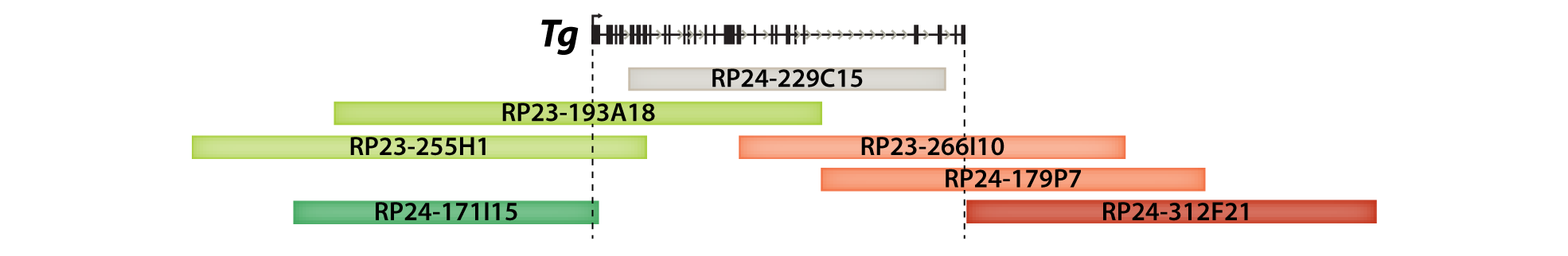 | |
| ***Ttn*** | |
| Gene body | RP23-310F9, RP23-22E10 |
| Gene halves | RP23-140P12, RP23-314B24 |
| Gene flanks | RP23-1M1, RP23-316G15 |
| 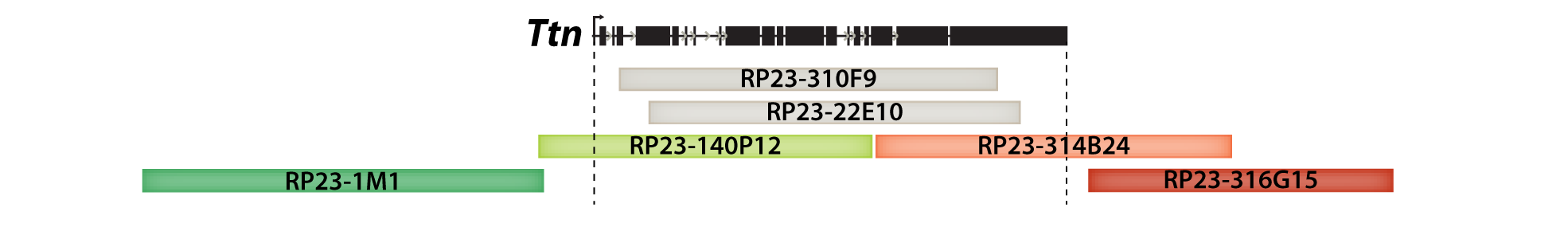 | |
| ***Neb*** | |
| Gene body | RP23-82J15 |
| Gene halves | RP23-91F4, RP23-177B19 |
| 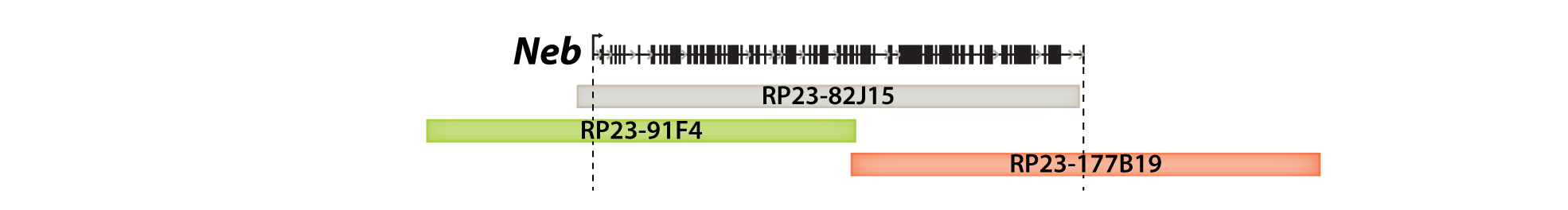 | |
| ***Myh11*** | |
| Gene body | RP24-158B9, RP23-465H10 |
| Gene halves | RP24-283P15, RP23-291J21 |
| Gene flanks | RP23-415B16, RP23-339L21 |
| 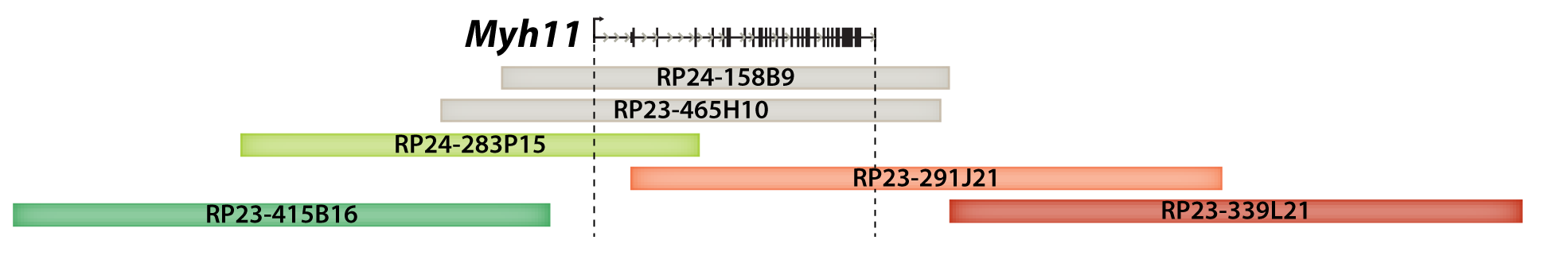 | |
| ***Cald1*** | |
| Gene body | RP23-265A17 + RP23-386A17 |
| 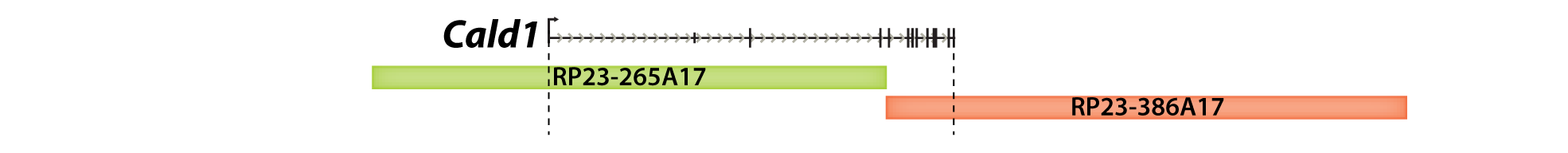 | |
